# Development of a cell-permeable Biotin-HaloTag ligand to explore functional differences between protein variants across cellular generations

**DOI:** 10.1101/2024.09.18.613519

**Authors:** Anoop Kumar Yadav, Abhijeet S. Jadhav, Pawel Szczepanik, Paolo Fagherazzi, Ivo Kabelka, Robert Vácha, Jakub Svenda, Hana Polasek-Sedlackova

## Abstract

HaloTag technology represents a versatile tool for studying proteins. Fluorescent HaloTag ligands employed in sequential labeling led to the discovery of distinct protein variants for histones, cohesins, and MCM complexes. Nonetheless, an efficient biochemical approach to separate the distinct protein variants to study their biological functions is missing. Principally being a gap in technology, the HaloTag toolbox lacks affinity ligands displaying good cell permeability and efficient affinity capture. Here, we describe the design, synthesis, and validation of a new cell-permeable Biotin-HaloTag ligand, which allows rapid labeling of Halo-tagged proteins in live cells and their efficient separation using streptavidin pull-down. Our work outlines how to use the herein-developed affinity ligand in sequential labeling to biochemically separate distinct protein variants and study their biological properties. The approach holds immense potential for addressing fundamental questions concerning essential cellular processes, including genome duplication and chromatin maintenance.

## INTRODUCTION

Proteins serve as the primary driving force behind nearly all cellular processes, and their proper functioning is essential for the health and fitness of living organisms. Therefore, exploring protein function represents a fundamental approach to unveiling the molecular nature of vital cellular processes and their related changes underlying disease development. A diverse array of biochemical, cell biology, proteomics, and genetics methodologies have been devised to visualize proteins and study their function through the attachment of a functional tag to the protein of interest. Over the years, a wide range of functional tags has been developed to suit different types of experimental approaches.

For instance, the polyhistidine tag (His-tag) is one of the most popular small tags for protein isolation and purification, while green fluorescent protein (GFP) and its derivatives are widely utilized in imaging studies^1,2^. More recently, small inducible tags, such as auxin-inducible degron (AID), have emerged as prominent tools for achieving rapid protein degradation in functional studies^3-5^. Nevertheless, a comprehensive understanding of protein function typically requires the combination of multiple tailor-made tags, which is a major limitation of the traditional protein tagging approach. The elegant solution for this bottleneck came with the discovery of self-labeling protein tags, exemplified by the HaloTag and SNAP-tag^6,7^. These tags, fused to the protein of interest, are based on the covalent interaction of an enzyme with a small synthetic ligand comprising a reactive group and a functional reporter group. While the SNAP-tag is the mutant of the DNA repair protein O6-alkylguanine-DNA alkyltransferase enzyme reacting with benzylguanine derivatives, the HaloTag is the mutant of bacterial haloalkane dehalogenase enzyme binding the reactive linear chloroalkanes^6,7^. Both enzymes react specifically and rapidly with their respective small-molecule ligands, leading to irreversible covalent labeling of HaloTag or SNAP-tag. A wide variety of functional reporter groups can be linked to the enzyme-reactive group of the ligands, lending the much-needed versatility to protein tagging. Nowadays, the commercially available HaloTag or SNAP-tag systems include a wide spectrum of fluorescent dyes for in vitro and in vivo imaging studies as well as conjugation of specific protein fragments, e.g., E3 ubiquitin ligase, enabling proximity-driven degradation of Halo-tagged proteins^8-12^. The operational simplicity of working with fully assembled bifunctional ligands may offer distinct experimental advantages over alternative two-step labeling approaches based on click chemistry^13^.

All these advances highlight the Halo and SNAP tags as powerful tools for studying protein function using complex biological approaches. The unique feature of these systems is the option for sequential labeling of proteins with different ligands using a pulse-chase approach to monitor protein dynamics, allowing for the real-time tracking of protein trafficking, synthesis, and overall turnover^14,15^. In particular, this method has emerged as a powerful approach for studying the maintenance of chromatin landscape through histone variants, one of the most prominent carriers of epigenetic memory necessary for preserving cellular identity over generations of dividing cells^16^. The Halo/SNAP-tag-based imaging systems have been successfully used to distinguish between old and new histones and visualize their dynamics throughout the cell cycle at the single-cell level^14,15^. The discoveries of various dedicated histone chaperones guiding the timing and mode of histone deposition during vital cellular processes, such as DNA replication, repair, and transcription, represent a major advance in our understanding of the molecular pathways responsible for chromatin landscape maintenance^17-20^. Beyond histones, distinct protein variants have been observed through HaloTag-based imaging within cohesin and, more recently, minichromosome maintenance (MCM) protein complexes^21-23^. Cohesin protein complexes play a pivotal role in maintaining the chromatin architecture by organizing the genome into dynamic chromatin loops and facilitating the cohesion of sister chromatids generated during genome duplication^24^. Recent research has delineated two discrete forms of cohesin protein complexes^21^. One variant is linked with chromatin post-mitotic exit and subsequently transforms into cohesive forms behind the replication forks during the S phase, while the other variant is newly loaded onto nascent DNA at the replication fork. Furthermore, the transition and de novo loading of cohesin complexes are facilitated by a distinct set of proteins reminiscent of histone dynamics^25^. MCM protein complexes are essential precursors of genome duplication^26^. During the G1 phase, a massive amount of MCM complexes is loaded on chromatin, the portion of which is converted to active replicative helicase, unwinding duplex DNA, during DNA replication in the S phase. Using the Halo-tag imaging system, our recent work revealed that MCM complexes exist in different protein forms–parental and nascent MCMs^23^. These protein forms serve distinct functions during the DNA replication program and are maintained by specific pathways with dedicated chaperones.

Despite the considerable progress in understanding the functional differences among protein variants of histone, cohesin, and MCM complexes, several fundamental questions remain unexplored. For instance, what determines the functional properties of different protein forms, and what chaperones and molecular pathways are involved in generating these variants? Additionally, can protein variants other than histones carry epigenetic information through cellular generations to sustain regulatory settings of chromatin and DNA replication? The experimental approach and robust technology enabling the efficient labeling (maximum labeling of a given protein pool within short time periods) of different protein variants in the cellular environment and their subsequent capture by affinity methods to answer these questions are currently not well developed. HaloTag or Snap-tag probes based on biotin-streptavidin interaction represent a logical extension of the self-labeling protein tag technology. For studies of distinct protein variants in the cell, such probes need to display good live-cell permeability, rapid protein labeling, and efficient affinity capture. Meeting these requirements has been challenging thus far, as highlighted by recent studies of the various Biotin-HaloTag constructs^27,28^. In this manuscript, we present the development and comprehensive characterization of a new cell-permeable Biotin-HaloTag ligand, which allows for efficient labeling of Halo-tagged proteins in living cells and biochemical separation of the protein variants using streptavidin pulldown.

## RESULTS

### The lack of suitable HaloTag ligands for efficient labeling and affinity capture of Halo-tagged proteins

Various beads and resins have been developed to capture and purify Halo- or SNAP-tagged proteins^29,30^. However, these tools are designed to interact with Halo or SNAP tags irrespective of the occupancy of the active site by specific ligands, thereby lacking the capability to separate distinct protein variants. In addition, new and old protein variants of histones, cohesins, and MCMs are generated in a cell-cycle-dependent manner^16,22,23^. Therefore, a maximum labeling efficiency of a given protein pool in live cells within a sufficiently short period of time (≤ 2 hours in the context of the 24-hour-long cell cycle) is the critical criterion when considering the experimental setup. Recognizing these limitations, we set out to explore a HaloTag-based experimental approach using biotin-streptavidin interaction for efficient labeling and affinity capture of Halo-tagged proteins (Figure 1A). Since two biotin-containing HaloTag ligands are commercially available (Figure 1B), namely HaloTag Biotin ligand (here referred to as Biotin-[0]-HaloTag ligand 1, or shortly Ligand 1) and HaloTag PEG-Biotin ligand (here referred to as Biotin-[16]-HaloTag ligand 2, or shortly Ligand 2), we first evaluated whether they meet the required properties outlined above. To this end, we employed human U2OS cells expressing endogenously Halo-tagged MCM4 subunit using CRISPR-Cas9 genome editing. The MCM4-Halo cell line was thoroughly tested to validate the homozygous tagging of all MCM4 alleles (Figures S1A-C). To test the labeling efficiency of commercial Biotin-HaloTag ligands, the MCM4-Halo cells were incubated with respective ligands at a final labeling concentration of 2.5 μM for 2 hours, followed by a collection of cell lysates and detection of labeled MCM4 by western blotting (Figure 1C). While Ligand 1 effectively labeled MCM4-Halo, Ligand 2 showed only a modest capacity for labeling (approximately 20-40%) of the endogenous MCM4 protein under the specified conditions (Figure 1C). To further determine whether Ligand 1 can efficiently label the entire available MCM4 protein pool present in the cells, we performed a pulse-chase labeling protocol (Figure 1D). First, cells were pulsed with Biotin-HaloTag ligands, and after a brief wash, fluorescence-based JFX554-HaloTag ligand was added after additional time elapsed. If Ligand 1 labels the entire MCM4 protein pool, then no fluorescence signal should be detected by the JFX554-HaloTag ligand. Indeed, the pulse-chase experiment revealed that Ligand 1 effectively labels the entire cellular fraction of MCM4 protein, while Ligand 2 shows only a modest labeling capacity (Figure 1D). The same observations were reproduced by orthogonal approach, during which the labeling efficiency of individual ligands was measured as a residual MCM fraction labeled by JFX554-HaloTag ligand at the single cell level using fully automated quantitative image-based cytometry (QIBC) (Figure 1E; Figure S1D). The labeling kinetics of both commercial ligands were additionally measured in a concentration- and time-dependent manner (Figures S1E, F). While Ligand 1 efficiently labeled almost the entire cellular fraction of MCM4-Halo at a final labeling concentration of 1 μM, the major fraction of MCM4 protein, approximately 50 %, remained unlabeled at the highest concentration of Ligand 2 (Figure S1E). QIBC of residual MCM4 fraction visualized by fluorescent ligand revealed that extending the treatment time to six hours is not sufficient to label the entire MCM4 fraction by Ligand 2 (Figure S1F). The observed slow kinetics of HaloTag labeling by Ligand 2 in the context of the 24-hour doubling time for the majority of human cells disqualifies this ligand for the purpose of affinity capture of different protein variants.

**Figure 1:**
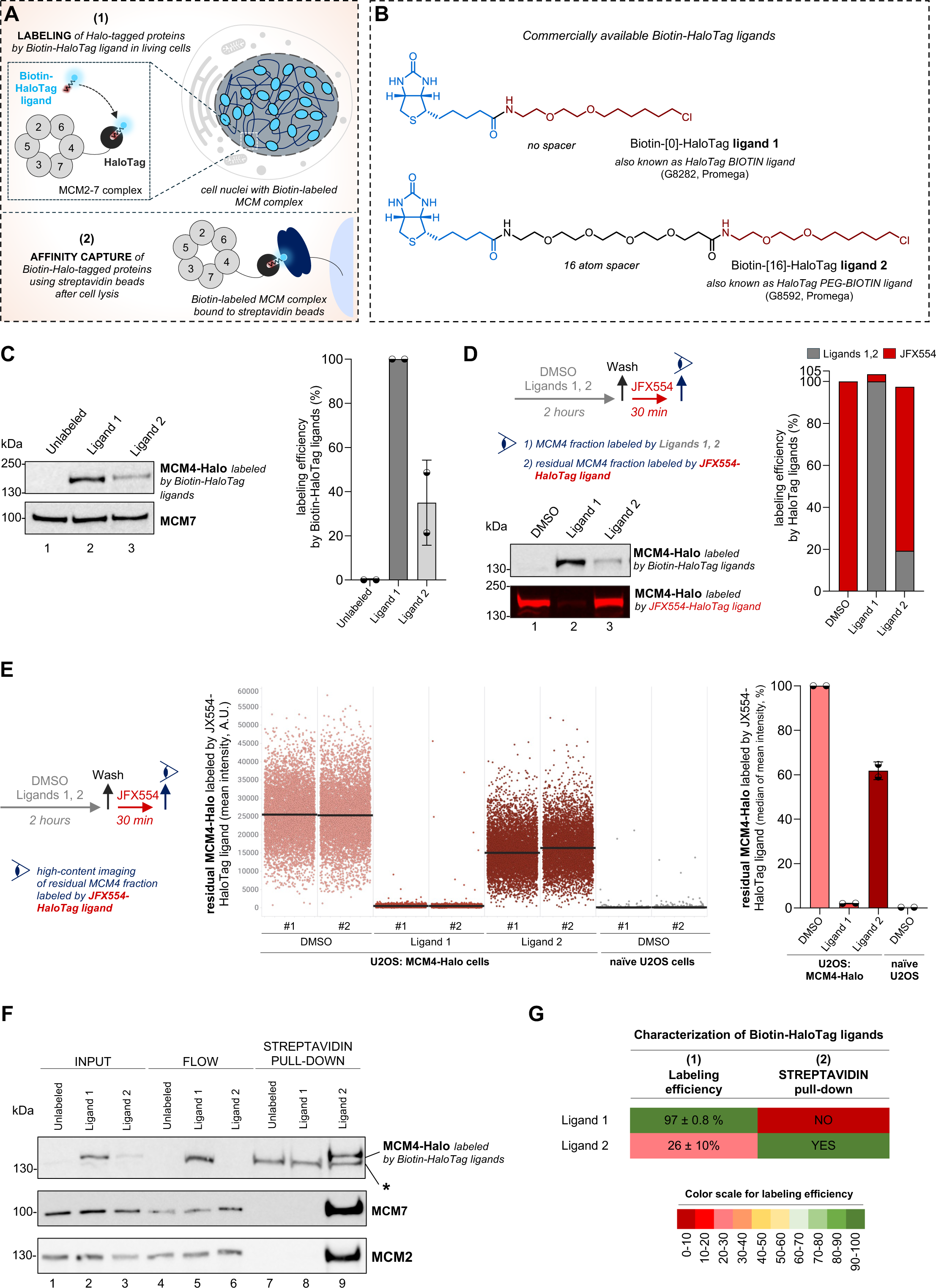
Commercially available Biotin-HaloTag ligands lack the necessary properties to discriminate between distinct protein variants. **(A)** A model describing a HaloTag experimental approach based on biotin-streptavidin interaction enabling efficient labeling protein variants (represented by MCM complex) in the cellular environment and subsequent affinity capture of Biotin-labeled proteins by streptavidin beads (see text for details). **(B)** Chemical structure of commercially available Biotin-HaloTag ligands. **(C)** Left, western blotting of whole cell lysates of MCM4-Halo U2OS cells labeled with the indicated Biotin-HaloTag ligands at a final concentration of 2.5 μM for 2 hours. MCM7 was stained as a loading control. Right, quantification of labeling efficiency for indicated Biotin-HaloTag ligands based on western blot on left. **(D)** Top left, the pulse-chase protocol of MCM4-Halo U2OS cells labeled with the indicated HaloTag ligands. Bottom left, SDS-PAGE or western blotting of whole cell lysates of MCM4-Halo U2OS cells labeled with indicated HaloTag ligands. Right, quantification of labeling efficiency for indicated Biotin-HaloTag ligands based on SDS-PAGE and western blot on left. **(E)** Left, the pulse-chase protocol of MCM4-Halo U2OS cells labeled with the indicated HaloTag ligands. Middle, QIBC of the residual fraction of MCM4-Halo labeled by JFX554-HaloTag ligand. Nuclear DNA was counterstained with DAPI. Lines denote medians; *n* ≈ 6000 cells per condition. Right, the quantification of QIBC plots in the middle. Each bar indicates the median of mean intensity normalized with respect to DMSO as 100 percent; *n* = 2 technical replicates. **(F)** Streptavidin pull-down of whole cell lysates of MCM4-Halo U2OS cells labeled with indicated Biotin-HaloTag ligands at a final concentration of 2.5 μM for 2 hours. The asterisk indicates an unspecific band from streptavidin agarose beads; see uncropped blots in Supplementary Information File 1. **(G)** Graphical summary of tested properties for commercial Biotin-HaloTag ligands. The labeling efficiency for individual ligands at a final concentration of 2.5 μM for 2 hours is presented as a mean with standard deviation based on western blotting and QIBC experiments in Figure 1 and S1.

The second criterion in our assessment of the two commercial Biotin-HaloTag ligands is their ability to engage with streptavidin beads. Such interaction enables the capture of biotin-labeled protein complexes, which can be further subjected to proteomic analysis to identify interacting partners or specific posttranslational modifications. Strikingly, a streptavidin-based pull-down experiment revealed that MCM4-Halo labeled by commercial Ligand 1 was unable to engage with streptavidin beads, as the entire pool of labeled protein was present in the flowthrough fraction (lane 5; Figure 1F). In sharp contrast to Ligand 1, Ligand 2, despite its reduced labeling efficiency (compare input fractions, lanes 2 and 3), was able to interact with streptavidin beads. Pulling down the MCM2 and MCM7 subunits along with MCM4-Halo validates the efficacy of this approach in capturing functional MCM complexes (Figure 1F). Taken together, our comprehensive characterization of Biotin-HaloTag ligands revealed that none of the currently commercially available ligands possess the necessary properties to biochemically study distinct protein variants (Figure 1G).

### Development of new Biotin-HaloTag ligands using variable-length atom spacers

Since commercial Ligand 1 failed in the streptavidin pull-down, we explored several strategies to enhance the poor labeling capacity of Ligand 2. We argued that the poor labeling of HaloTag by Ligand 2 may be attributed to low cell permeability^28^. To investigate this aspect, we employed verapamil, a broad-spectrum efflux pump inhibitor previously shown to enhance the labeling efficiency of fluorogenic probes for live cell imaging of the cytoskeleton^31^. Pulse-chase experiments, analyzed either by western blotting (Figure 2A) or QIBC (Figure 2B, Figure S2A), indeed demonstrated an increase in the labeling efficiency of Ligand 2 in the presence of verapamil. This supports the notion that the poor labeling efficiency is attributable to the ligand’s impaired cell permeability rather than its degradation within the cellular environment. However, it is important to note that while verapamil can be advantageous for short-term experiments, it may not be suitable for long-term studies, such as studying different protein forms produced in two successive cellular generations. As shown previously and in this study, the use of verapamil can affect cell growth (Figure 2C; Figures S2B, C)^32^.

**Figure 2:**
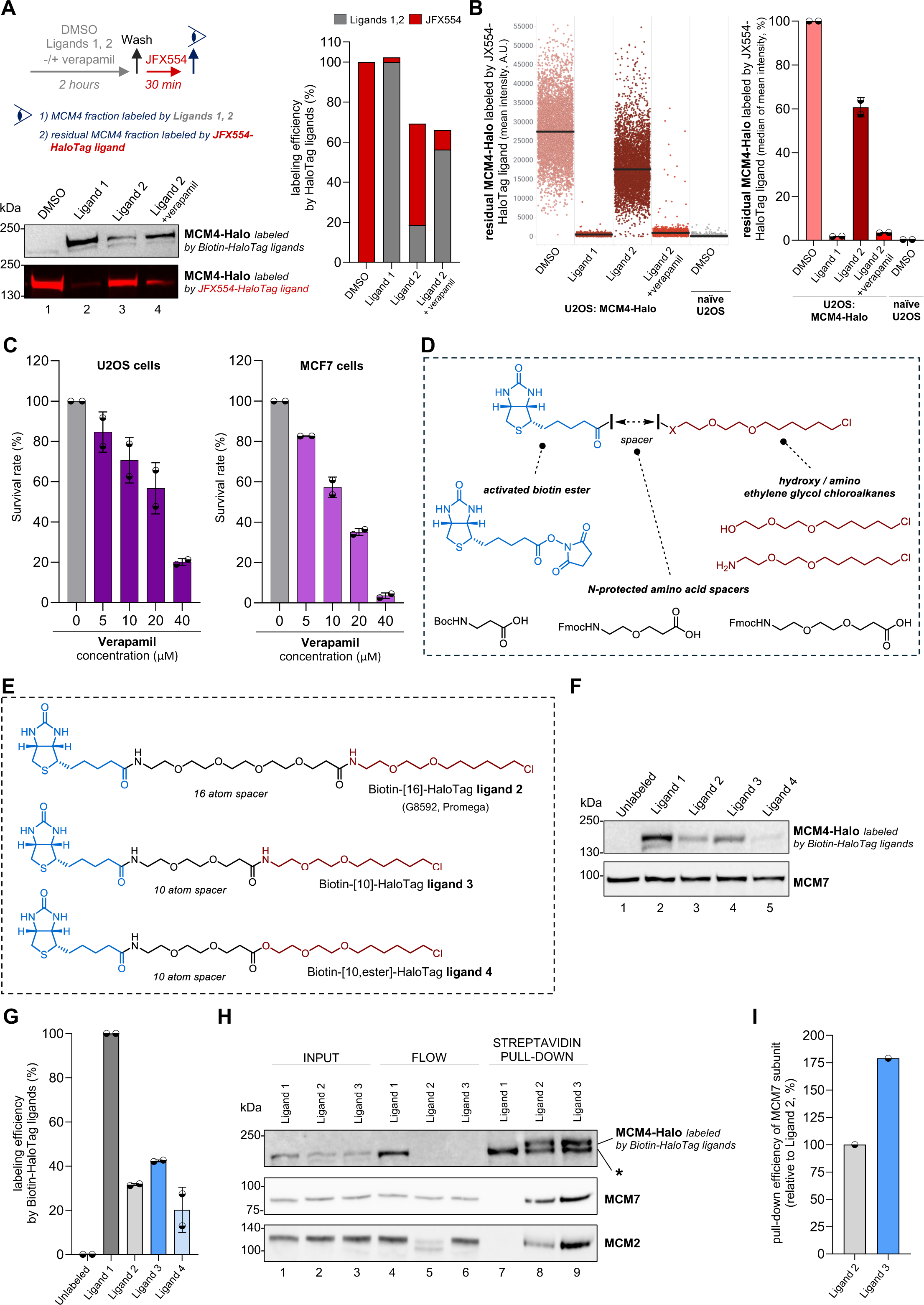
Development of new Biotin-HaloTag ligands by modifying chemical properties of amino acid spacers. **(A)** Top left, the pulse-chase protocol of MCM4-Halo U2OS cells labeled with indicated HaloTag ligands along with or without 10 μM verapamil. Bottom left, SDS-PAGE or western blotting of whole cell lysates of MCM4-Halo U2OS cells labeled with indicated HaloTag ligands. Right, quantification of labeling efficiency for indicated Biotin-HaloTag ligands based on SDS-PAGE and western blot on left. **(B)** Left, QIBC of the residual fraction of MCM4-Halo labeled by JFX554-HaloTag ligand. Nuclear DNA was counterstained with DAPI. Lines denote medians; *n* ≈ 5500 cells per condition. Right, the quantification of QIBC plots on the left. Each bar indicates the median of mean intensity normalized with respect to DMSO as 100 percent; *n* = 2 technical replicates. **(C)** Clonogenic survival of U2OS (left) and MCF7 (right) cells after treatment with increasing doses of verapamil. Each bar indicates the average of observed colonies normalized with respect to untreated cells as 100 percent; *n* = 2 biological replicates. **(D)** Principal fragments used for the synthesis of all Biotin-HaloTag ligands studied in this work (see Supplementary Information File 2 for detailed synthetic procedures). **(E)** Chemical structures of the commercially available Biotin-HaloTag ligand 2 with 16-atom spacer (top) and synthesized Biotin-HaloTag ligands 3 and 4 with 10-atom spacers (middle, bottom). **(F)** Western blotting of whole cell lysates of MCM4-Halo U2OS cells labeled with indicated Biotin-HaloTag ligands at a final concentration of 2.5 μM for 2 hours. MCM7 was stained as a processing control. **(G)** Quantification of labeling efficiency for indicated Biotin-HaloTag ligands based on western blot in (F). **(H)** Streptavidin pull-down of whole cell lysates of MCM4-Halo U2OS cells labeled with indicated Biotin-HaloTag ligands at a final concentration of 2.5 μM for 2 hours. **(I)** Quantification of pull-down efficiency of MCM7 subunit by the indicated Biotin-HaloTag ligands based on western blot in (H).

In order to develop a new Biotin-based HaloTag ligand capable of efficient labeling and streptavidin pull-down with minimal cell toxicity, we sought to modify the structure of commercial Biotin-HaloTag ligands (Figure 1B). We hypothesized that the inability of the cell-permeable Ligand 1 to bind the streptavidin beads after its covalent attachment to HaloTag protein might be due to an insufficient atom spacer between the biotin group and the reactive chloroalkane tail. The analysis of available crystal structures revealed that the chloroalkane chain is buried deep within the HaloTag active site. For Ligand 1, this could render the biotin group sterically inaccessible and compromise the interaction with streptavidin. Conversely, the 16-atom spacer in Ligand 2 allows ample space between the biotin and chloroalkane groups necessary for simultaneous interaction with HaloTag and streptavidin, however, at the expense of low labeling efficiency due to presumed poor cell permeability (see above). Following the rationale, we used chemical synthesis to prepare a set of Biotin-HaloTag ligands featuring variable-length amino acid spacers, which, we hoped, could combine the desired characteristics of the two commercial ligands. Our synthetic approach is overviewed in Figure 2D. Inspired by prior literature^7,33^, the known amine-substituted chloroalkane component was coupled to various *N*-protected amino acid spacers. Following the removal of the nitrogen-protecting group, biotin was attached via its active (NHS) ester. Using the outlined synthetic approach, we first produced two Biotin-HaloTag ligands denoted as Ligands 3 and 4, each containing a 10-atom spacer (Figure 2E). Ligand 4, prepared from the hydroxy-substituted chloroalkane component, is an ester analog of the known Ligand 3(ref.^33^) and was chosen in light of the recent report on amide-to-ester substitution as a strategy to increase cell permeability of PROTAC constructs^34^. The labeling efficiency of Ligands 3 and 4 was examined by western blotting (Figures 2F, G). While Ligand 3 showed slightly increased labeling efficiency, its ester analog (Ligand 4) was inferior relative to the commercial Ligand 2. The modest increase in labeling efficiency seen with Ligand 3 suggested that shortening the atom spacer may be used to improve the labeling properties. Next, to investigate whether Ligand 3, having the shorter atom spacer (relative to Ligand 2), retains its ability to bind streptavidin, we performed the streptavidin pull-down (Figures 2H, I). The results confirmed that Ligand 3-HaloTag conjugate maintained the capacity to interact with streptavidin beads. The functional relevance of this interaction was demonstrated by the pull-down of other MCM subunits, specifically MCM7 and MCM2, indicating the successful capture of functional MCM complexes. Overall, these findings suggested that a judicious design of the atom spacer in the Biotin-HaloTag ligands is a viable strategy to enhance labeling capacity while preserving the ability to interact with streptavidin beads.

### Identification and characterization of new Biotin-HaloTag ligand with efficient labeling in living cells and streptavidin pull-down capacity

Motivated by observations in Figure 2, we decided to shorten the 10-atom spacer further to attain the labeling efficiency of commercial Ligand 1. To help determine the appropriate spacer length, we created computational models for visualization of the HaloTag-ligand-streptavidin ternary complexes. These were assembled from the available crystal structures of HaloTag-chloroalkane and streptavidin-biotin binary complexes, respectively. First, an atom spacer of a chosen length was attached to the carboxylate of biotin within the streptavidin-biotin complex (PDB: 3RY2; a single subunit of streptavidin tetramer was used)^35^, and multiple conformers of the atom spacer were generated by constrained embedding in RDKit. Only the lowest energy spacer conformers (within 1 kcal/mol) were kept. Then, the HaloTag-chloroalkane complex (PDB: 6U32, modified)^9^ was attached to the carboxy group of each of the computed spacer conformations using the least squares fit of three atoms. Finally, the ternary complexes were ranked using the number of overlapping heavy atoms. The representative structural visualizations of the commercial ligands corroborate our earlier experimental findings (Figure 1). It is evident that Ligand 1 is too short to accommodate both HaloTag and streptavidin proteins (left, Figure 3A), in contrast to Ligand 2 having a 16-atom spacer (right, Figure 3A) and Ligand 3 with a 10-atom spacer (Figure S2D). Based on the structural analysis, we designed a new ligand with a 7-atom spacer (Ligand 5, middle in Figure 3A). Despite being nine atoms shorter than commercial Ligand 2, Ligand 5 was predicted in silico to support the formation of a ternary HaloTag-ligand-streptavidin complex.

**Figure 3:**
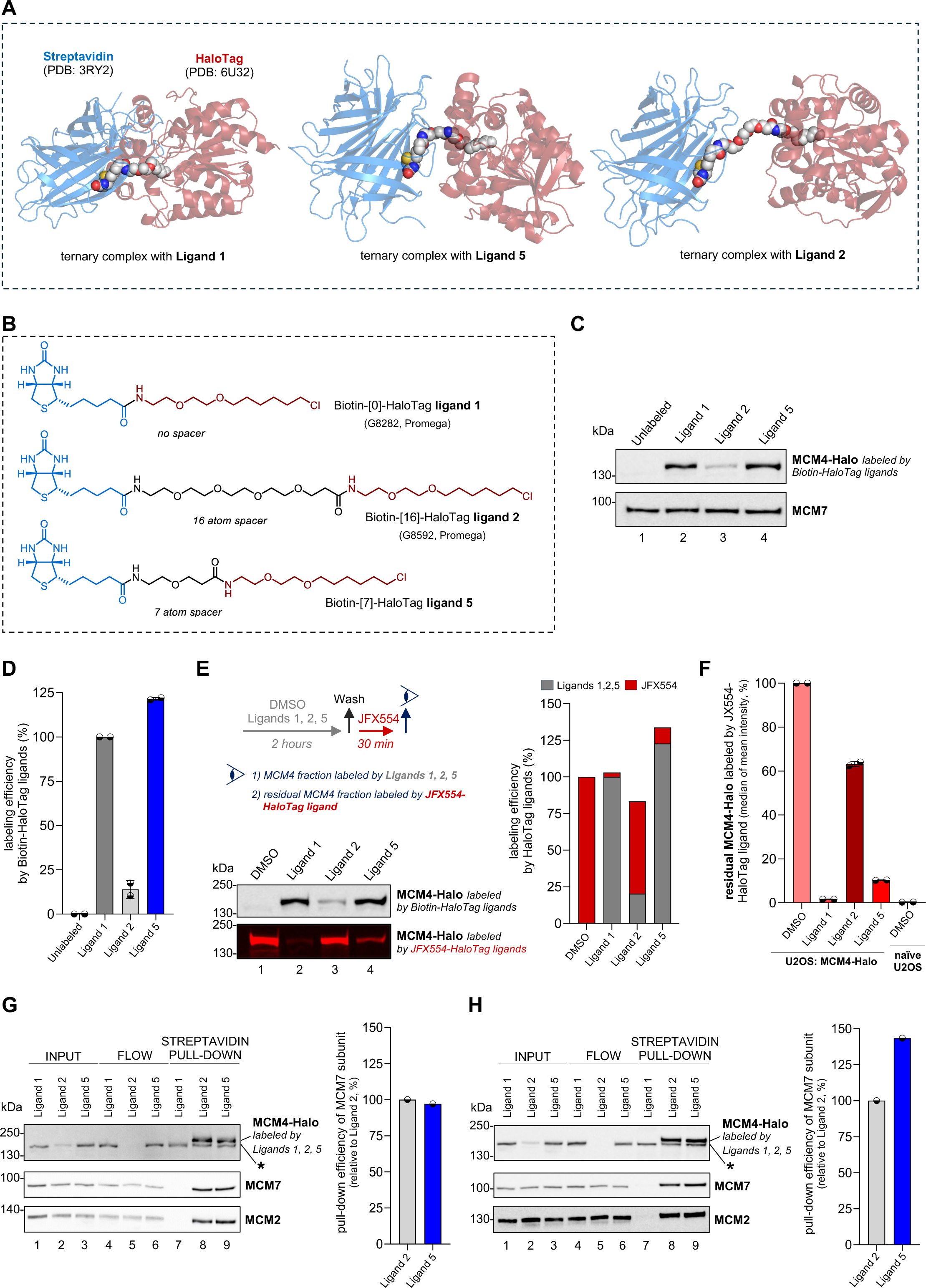
Biotin-[7]-HaloTag ligand 5 with 7-atom spacer is proficient in cell labeling assays and streptavidin pull-down experiments. **(A)** Representative structural visualizations of the HaloTag-ligand-streptavidin ternary complexes for ligands 1, 2, and 5. The complexes were calculated from the binary complexes of streptavidin-biotin (PDB: 3RY2, one subunit of the tetramer) and HaloTag-chloroalkane (PDB: 6U32, modified), with the exhaustive generation of spacer conformations by constrained embedding in RDKit. **(B)** Chemical structures of commercially available Biotin-HaloTag ligands 1 and 2 (top, middle) and newly designed Biotin-HaloTag ligand 5 with a 7-atom spacer (bottom). **(C)** Western blotting of whole cell lysates of MCM4-Halo U2OS cells labeled with indicated Biotin-HaloTag ligands at a final concentration of 2.5 μM for 2 hours. MCM7 was stained as a processing control. **(D)** Quantification of labeling efficiency for indicated Biotin-HaloTag ligands based on western blot in (B). **(E)** Top left, the pulse-chase protocol of MCM4-Halo U2OS cells labeled with the indicated HaloTag ligands. Bottom left, SDS-PAGE or western blotting of whole cell lysates of MCM4-Halo U2OS cells labeled with indicated HaloTag ligands. Right, quantification of labeling efficiency for indicated Biotin-HaloTag ligands based on SDS-PAGE and western blot on left. **(F)** The quantification of the residual fraction of MCM4-Halo labeled by JFX554-HaloTag ligand. Each bar indicates the median of mean intensity normalized with respect to DMSO as 100 percent; *n* = 2 technical replicates. See QIBC in Supplementary Figure 2D. **(G)** Left, Streptavidin pull-down of whole cell lysates of MCM4-Halo U2OS cells labeled with indicated Biotin-HaloTag ligands at a final concentration of 2.5 μM for 2 hours. Right, quantification of pull-down efficiency of MCM7 subunit by the indicated Biotin-HaloTag ligands based on western blot on the left. **(H)** Left, Streptavidin pull-down of whole cell lysates of MCM4-Halo U2OS cells labeled with indicated Biotin-HaloTag ligands at a final concentration of 2.5 μM for 2 hours. Right, quantification of pull-down efficiency of MCM7 subunit by the indicated Biotin-HaloTag ligands based on western blot on the left.

Following our synthetic scheme (Figure 2D), we prepared the above-designed Ligand 5 (Figure 3B). Notably, western blot analysis demonstrated that Ligand 5 achieves high labeling of Halo-tagged MCM4 comparable to that of commercial Ligand 1 (Figures 3C, D), demonstrating that shortening the spacer from 16 to 7 atoms substantially enhanced the labeling efficiency. In addition, the labeling efficiency was tested by pulse-chase experiments and evaluated by western blotting and QIBC, indicating that Ligand 5 proficiently labels nearly the entire fraction of MCM4 protein in the cellular environment (Figures 3E, F; Figures S2E, F). Furthermore, comparisons of the labeling kinetics of Ligand 5 and commercial Ligand 1 in a concentration-dependent manner revealed that, despite slower kinetics, Ligand 5 achieved approximately 90% labeling of the cellular MCM4 fraction at higher ligand concentrations, as corroborated by both western blotting and QIBC analyses (Figures S3A, B). The robustness of our synthesis and data was confirmed in experiments assessing the labeling kinetics in a concentration- and time-dependent manner using two independent batches of the synthetic Ligand 5 (Figures S3C, D). Both batches of Ligand 5 showed the same labeling kinetics, achieving approximately 90% labeling of the cellular MCM4 fraction at a concentration of 2.5 μM within two hours. Altogether, our findings unequivocally demonstrate that the introduction of a 7-atom spacer into the Biotin-HaloTag ligand substantially increased the labeling efficiency of Halo-tagged proteins.

As Ligand 5 demonstrates a comparable labeling efficiency to commercial Ligand 1, we next evaluated the newly synthesized ligand for its ability to interact with streptavidin beads as predicted by our structural visualizations. This was done using a streptavidin-based pull-down assay (Figure 3G), which demonstrated that Ligand 5 in conjugation with HaloTag retains its ability to bind streptavidin. Importantly, interaction with other MCM subunits was observed, indicating the successful capture of functional MCM complexes through Ligand 5-HaloTag conjugation. Although the efficiency of streptavidin pull-down mirrored that of commercial Ligand 2, a substantial portion of MCM4-Halo labeled by Ligand 5 was evident in the flowthrough fraction (compare lanes 5 and 6 of the western blot in Figure 3G). This observation may be rationalized by the approximately 5-fold higher labeling efficiency of Ligand 5 compared to commercial Ligand 2 (for further details, see quantification in Figure 3D, or compare lanes 2 and 3 in the input fraction of Figure 3G). Such increased labeling efficiency could potentially exhaust the pull-down capacity of the streptavidin beads. In support of this argument, upon increasing the amount of streptavidin beads, we enhanced the pull-down efficiency for MCM4-Halo labeled by Ligand 5 (compare Figures 3G and 3H). Noteworthy, the efficient binding to streptavidin can be utilized beyond affinity pull down, for instance, the visualization of protein complexes directly in their natural environment. Thus, we employed Alexa Fluor 647-conjugated streptavidin to detect Ligand 5-bound MCM-Halo complexes within the cell nucleus using QIBC. The analysis confirmed the capacity of Ligand 5 to label nuclear as well as chromatin-bound fractions of MCM complexes (Figures S4A, B). No signal was observed in naïve U2OS cells, indicating that labeling by Ligand 5 is HaloTag-specific. Additionally, in alignment with previous literature^36^, typical chromatin dynamics of MCM complexes labeled by Ligand 5 were evident throughout the cell cycle, indicating the visualization of MCM complexes involved in the genome duplication process (Figures S4B, C).

The successful generation and favorable properties of Biotin-HaloTag ligand 5 prompted us to examine an analogous ligand having only a 4-atom spacer (Ligand 6, Figure S4D). The structural visualization model introduced above (Figure 3A; Figure S4E) suggested the 4-atom spacer to be a borderline case and had to be assessed experimentally. The synthesis of Ligand 6 was straightforward following our approach (Figure 2D), as the spacer came in the form of commercially available *N*-protected β-alanine. While Ligand 6 showed marginal improvement in the labeling efficiency of Halo-tagged MCM4 compared to Ligand 5, it lost the ability to interact with streptavidin (i.e., ternary complex formation; Figures S4F, G). These observations underscore the importance of judicious optimization of the atom spacer in the design of Biotin-HaloTag ligands. The newly identified Ligand 5 with a 7-atom spacer is proficient in cell labeling assays and streptavidin pull-down experiments and superior to the commercially available Biotin-HaloTag ligands. These properties render Ligand 5 a candidate small-molecule tool for studying the biochemical properties of distinct protein variants.

### Experimental design to explore biochemical properties of protein variants represented by MCM and histone complexes

As mentioned in the preceding paragraphs, studying different protein forms across successive cellular generations may require longer exposure of cells to the new Biotin-HaloTag ligand 5. Therefore, we determined the effects of Ligand 5 on cell cycle progression and growth using QIBC and cell survival assays. The results demonstrated that extended exposure to Ligand 5 did not elicit alterations in the cell cycle (Figures S5A, B), nor did it impede the growth of U2OS and MCF7 cells (Figure 4A; Figures S5C, D). These findings, along with prior characterizations, suggest that Ligand 5 is well-suited for long-term studies, such as investigating the various protein forms generated across multiple cell divisions (Figure 4B).

**Figure 4:**
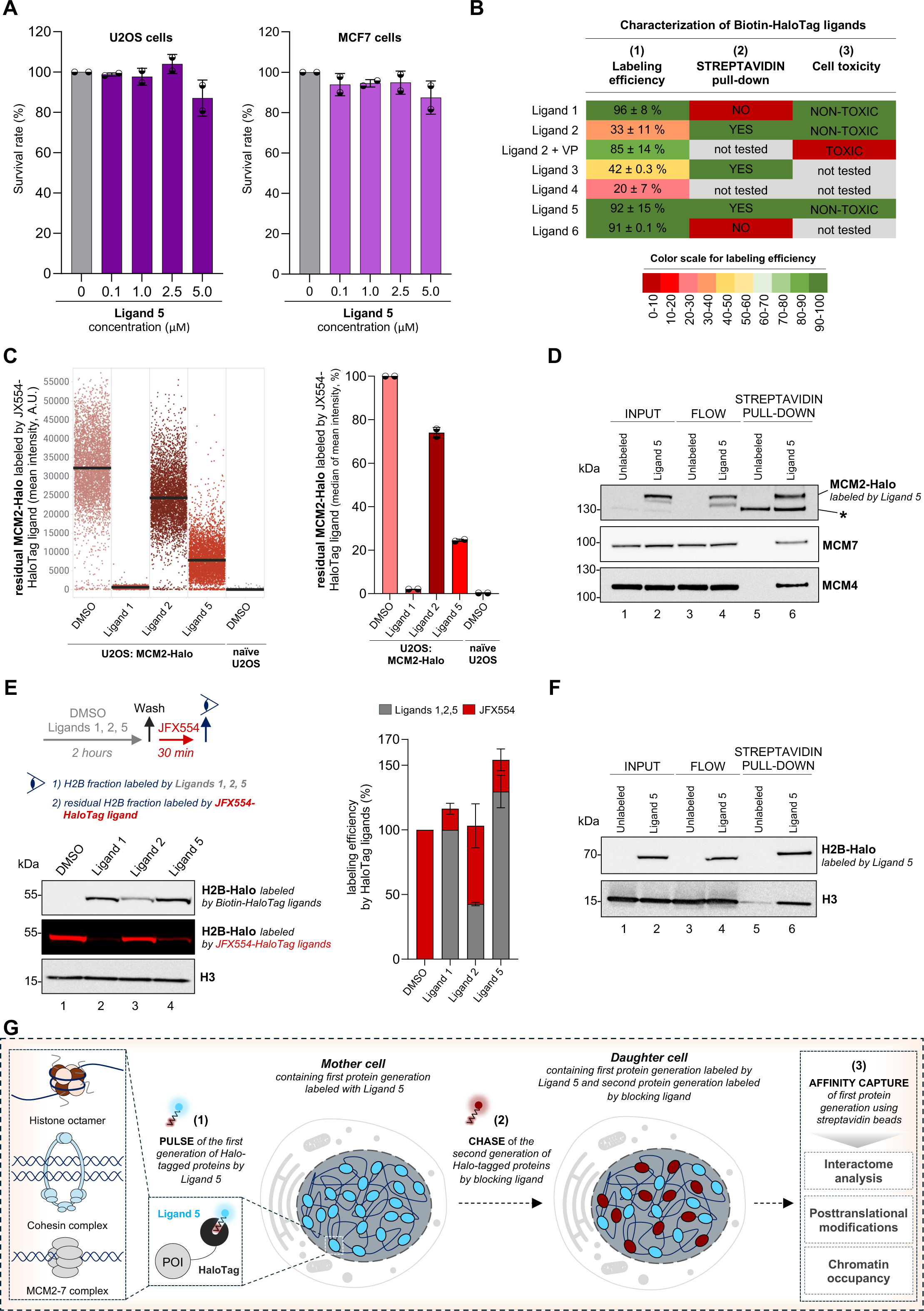
Ligand 5 represents a powerful tool to biochemically separate different protein variants. **(A)** Clonogenic survival of U2OS (left) and MCF7 (right) after treatment with increasing doses of Ligand 5. Each bar indicates the average of observed colonies normalized with respect to untreated cells as 100 percent; *n* = 2 biological replicates. **(B)** Graphical summary of tested properties for commercial and newly generated Biotin-HaloTag ligands. The labeling efficiency for individual ligands at a final concentration of 2.5 μM for 2 hours is presented as a mean with standard deviation based on all western blotting and QIBC experiments performed in this work. VP denotes verapamil. **(C)** Left, QIBC of the residual fraction of MCM2-Halo labeled by JFX554-HaloTag ligand. Nuclear DNA was counterstained with DAPI. Lines denote medians; *n* ≈ 5000 cells per condition. Right, the quantification of QIBC plots on the left. Each bar indicates the median of mean intensity normalized with respect to DMSO as 100 percent; *n* = 2 technical replicates. **(D)** Streptavidin pull-down of whole cell lysates of MCM2-Halo U2OS cells labeled with Ligand 5 at a final concentration of 2.5 μM for 2 hours. **(E)** Top left, the pulse-chase protocol of H2B-Halo HeLa cells labeled with indicated HaloTag ligands. Bottom left, SDS-PAGE or western blotting of whole cell lysates of H2B-Halo HeLa cells labeled with indicated HaloTag ligands. Right, quantification of labeling efficiency for indicated Biotin-HaloTag ligands based on SDS-PAGE and western blot on the left. **(F)** Streptavidin pull-down of whole cell lysates of H2B-Halo HeLa cells labeled with Ligand 5 at a final concentration of 2.5 μM for 2 hours. **(G)** A model describing a HaloTag experimental approach for studying biological properties of different protein variants through their biochemical separation based on biotin-streptavidin interaction. Previously generated cell lines and described pulse-chase protocols to distinguish protein variants of histones, cohesins, and MCM complexes can be modified in order to biochemically capture and separate these variants produced during two cellular generations. For specific biochemical separation of old (parental) protein variants, we recommend the usage of Ligand 5 in the pulse and blocking ligand during the chase. This will enable the labeling of the first protein generation by Ligand 5 and its separation from the second protein generation by streptavidin pull-down. For specific biochemical separation of new (nascent) protein variants, we suggest using a blocking ligand in the pulse and Ligand 5 during the chase. Any HaloTag ligand can serve as the blocking ligand. The described approach will enable the study of the respective interactomes, posttranslational modifications, and chromatin occupancy of different protein variants, thereby aiding in elucidating fundamental questions about genome duplication and chromatin maintenance (see text for details). POI represents the protein of interest.

Throughout the manuscript, we systematically evaluated the labeling efficiency and streptavidin pull-down capacity of commercial and newly synthesized Biotin-HaloTag ligands in U2OS cells, endogenously expressing the Halo-tagged MCM4 subunit. To reinforce our findings beyond a single protein target, we opted to broaden our exploration of Biotin-HaloTag ligands with different cell systems expressing distinct Halo-tagged proteins. To achieve this, we implemented CRISPR-Cas9 endogenous tagging to establish a U2OS cell line expressing Halo-tagged MCM2. The resulting cell line was extensively validated for the homozygous tagging of all MCM2 alleles (Figures S5E-H). Importantly, Ligand 5 exhibited high labeling efficiency of Halo-tagged MCM2 and demonstrated the capacity to effectively pull down functional MCM complexes using streptavidin beads (Figures 4C, D; Figure S5H). These results validate Ligand 5 as a tool for efficient labeling and biochemical separation of different protein variants within MCM complexes. Next, we employed human HeLa cells that ectopically expressed a Halo-tagged histone variant H2B, a commonly used tool for studying chromatin architecture^37^. The expression level of H2B-Halo in the cells was validated by western blotting and QIBC (Figures S6A, B). Notably, Ligand 5 exhibited a high labeling capacity for Halo-tagged H2B, well comparable to that of commercial Ligand 1, as measured by western blotting and QIBC (Figure 4E; Figures S6C, D). Moreover, using Ligand 5 in conjugation with Halo-tagged H2B, we successfully captured functional nucleosomes through streptavidin pull-down (Figure 4F). Collectively, we conclude that the Biotin-HaloTag ligand 5 is a new ligand encompassing efficient labeling and streptavidin pull-down capacity, properties that are unmatched by the commercially available ligands.

Based on our observations, we propose utilizing the new Ligand 5 in pulse-chase experiments to investigate the biological properties of different protein variants through their biochemical separation (Figure 4G). Previous studies have provided a detailed description of pulse-chase protocols to visualize different protein variants of Halo-tagged histones, cohesins, and MCM complexes generated through two successive cellular generations using one or two fluorescent HaloTag ligands^14,22,23^. We posit that the previously established cell lines and protocols can be readily adapted for the biochemical capture and isolation of diverse protein variants. For instance, Ligand 5 can be used to pulse the first protein generation, followed by a chase of the second protein generation by a blocking ligand, which can be any HaloTag ligand blocking the active site of the HaloTag. This experimental approach allows specific biotin-based labeling of old (parental) protein variants in daughter cells. Subsequently, these labeled proteins can be subjected to streptavidin-based affinity capture and purification to study specific biological properties of old (parental) protein variants. Following a similar logic, the exchange of Ligand 5 and the blocking ligand in pulse-chase experiments will enable the selective biochemical separation of new (nascent) protein variants. While this methodology holds promise for application to proteins other than histones, cohesins, and MCM complexes, it is important to note that optimization of the labeling intervals of pulse-chased protocols will be required to align with the cellular turnover of the protein of interest. In summary, we anticipate that the outlined experimental strategy can be employed to biochemically separate distinct protein variants, facilitating the investigation of their respective interactomes, posttranslational modifications, and chromatin occupancy using advanced proteomic and next-generation sequencing techniques. This, in turn, will provide mechanistic insights into fundamental cellular processes, including genome duplication, chromatin maintenance, and transmission of epigenetic information across generations of dividing cells. We elaborate on the specific biological questions that could be addressed using the Ligand 5-based approach in the last paragraphs of the discussion below.

## DISCUSSION

HaloTag technology represents a versatile platform for studying protein function using a range of biological methods^38^. Its versatility stems from the modular nature, and synthetic accessibility of HaloTag ligands, each incorporating different functional reporter groups conjugated to the reactive chloroalkane. Fluorescent HaloTag probes, particularly based on the series of rhodamine dyes, are now widely utilized in various biochemical and biological imaging studies^8,12,39,40^. A recently developed HaloPROTAC ligand, which facilitates the rapid degradation of Halo-tagged proteins via von Hippel Lindau (VHL) E3 ubiquitin ligase-based ligand, greatly expands the use of HaloTag technology beyond imaging studies^10^. However, a HaloTag ligand that would enable effective labeling of cellular proteins and their pull-down by affinity methods has been missing from the available HaloTag toolbox.

HaloTag affinity ligands based on the biotin-streptavidin interaction would represent a potentially powerful tool for the affinity capture and purification of Halo-tagged proteins and, in particular, for the biochemical separation of various protein variants generated in successive cellular generations. Currently, two biotin-based HaloTag ligands are available commercially (Ligand 1 and Ligand 2). Unfortunately, our comprehensive characterization of these two ligands revealed that they do not display optimal properties. While Ligand 1 achieves high labeling efficiency of the cellular fraction of Halo-tagged proteins, it fails in streptavidin pull-down experiments. Conversely, Ligand 2, after conjugation with HaloTag, effectively interacts with streptavidin but shows poor labeling efficiency of Halo-tagged proteins in the cellular environment due to limited cell permeability. These experiments, paired with computer-assisted structural visualizations performed, pointed to the critical nature of the atom spacer length situated between the biotin functional group and the reactive chloroalkane group. Our findings aligned well with a previous study, in which this issue was noted but not addressed experimentally^27^. The comprehensive characterization of the new ligands prepared in this work demonstrates how the judicious choice of the linker length can finetune the properties of the HaloTag ligands.

The newly identified Ligand 5 (Biotin-[7]-HaloTag ligand 5) effectively labels Halo-tagged proteins in the living cells and enables their pull-down through streptavidin beads without inducing cytotoxicity. We conducted extensive validation of the properties of Ligand 5 for various proteins and cell lines using a comprehensive array of techniques encompassing labeling, pulse-chase procedures, and streptavidin pull-downs, assessed via western blotting and high-content imaging. Although Ligand 5 exhibited slower kinetics in labeling Halo-tagged proteins in the cellular environment compared to commercial Ligand 1, it still efficiently labeled the entire protein pool of Halo-tagged proteins in a relatively short time (2 hours) and at low concentrations (2.5 μM). In sharp contrast to commercial Ligand 1, however, Ligand 5 can be readily captured by streptavidin beads. Based on these data, we believe that Ligand 5 effectively fills the gap in the palette of available HaloTag ligands for labeling and affinity pull-down of Halo-tagged proteins in their cellular environment. The tool empowers the HaloTag technology to investigate the functional differences among protein variants generated through two successive cellular generations (Figure 4G). In the past, Halo- or SNAP-tag-based pulse-chase experiments using fluorescent ligands across cellular generations have proven to be an invaluable method for uncovering distinct protein variants within histone, cohesin, and MCM complexes^14,22,23^. In the last paragraphs, we outline how the Ligand 5-based pulse-chase experiments can be used to capture and biochemically separate different protein pools produced by successive cellular generations and how this approach can help answer fundamental questions about the regulation of genome duplication and chromatin maintenance in the future.

Genome maintenance processes, such as DNA replication and repair, are vital for preserving the integrity of genetic information throughout cellular generations. Although essential, these processes disrupt the chromatin landscape, which is crucial for preserving the gene expression program that governs cell identity. The nucleosome, a histone octamer, constitutes the fundamental unit of chromatin architecture and is necessitated to be dismantled by an incoming replication fork to duplicate DNA and subsequently reassembled at the newly synthesized DNA^16^. More than forty years of extensive research have unveiled that old histones preceding the DNA replication fork are recycled within post-replicative chromatin alongside the deposition of newly synthesized histones. These old and new histone variants differ in molecular pathways responsible for their deposition, as well as through carried posttranslational modifications, representing a central source of cellular epigenetic memory^16^. Recent advancements in sequencing-based methodologies have furnished valuable insights into the molecular mechanisms underpinning the recycling of old histones at the replication fork^41-45^. In view of these recent breakthroughs, we hypothesize that employing Ligand 5-based pulse-chase experiments in combination with the iPOND (isolation of proteins on nascent DNA) method may serve as an intriguing approach to elucidate the interactomes of new and old histones at the replication fork (Figure 4G). Such an approach holds promise for unveiling novel histone chaperones and molecular mechanisms responsible for the maintenance of chromatin landscape and, thereby, the preservation of epigenetic memory.

Similar to histones, cohesin complexes play a crucial role in maintaining chromatin architecture, and their behavior during the cell cycle is reminiscent of that of histones. Initially, the cohesin complex was known for its function in holding sister chromatids during genome duplication; nevertheless, later on, its major role in organizing the topology of interphase chromatin was discovered^24^. These observations raised an important conundrum, namely, what governs the regulation of cohesin activity to transition from loop extrusion of pre-replicative chromatin to sister chromatid cohesion in post-replicative chromatin^21^. Previous research indicated that the answer may lie in understanding the dynamics of different forms of cohesin during the cell cycle. While one cohesin variant promotes loop extrusion after post-mitotic exit and, during the S phase, is transformed into cohesive forms behind the replication forks, the other variant is newly loaded onto nascent DNA at the replication fork^21^. Furthermore, recent studies revealed that transition and de novo loading of cohesin complexes are facilitated by a distinct set of proteins, reviewed in^25^. In light of these recent findings, we speculate that employing Ligand 5 in pulse-chase experiments in combination with the mass-spectrometry method may serve as a potent approach to explore the interactomes and posttranslational modifications of different cohesin variants (Figure 4G), thereby offering novel and valuable insights into the regulation of cohesin enzymatic activities on pre- and post-replicative chromatin.

Finally, our recent work has unveiled distinct protein variants within MCM complexes, the central players of genome duplication^23^. Upon inheritance, daughter cells receive both parental and nascent protein forms, each fulfilling specific functions during the DNA replication process, and their levels are preserved through distinct pathways. While parental MCMs are preferentially converted to active replisomes, the nascent MCMs remain largely inactive but serve as natural replisome pausing sites, sustaining physiological replication fork speed. Despite the valuable insights provided by our work into MCM biology, the key questions remained unresolved. Specifically, what defines the functional properties of parental and nascent MCMs that govern their functions during the DNA replication process? Furthermore, what protein chaperones are responsible for MCM biogenesis and recycling pathways? Building on our findings in this work, we propose that combining Ligand 5-based pulse-chase experiments with mass-spectrometry may represent an effective strategy to explore the posttranslational modifications of different MCM protein variants and identify novel protein chaperones crucial for maintaining MCM equilibrium (Figure 4G). Furthermore, the integration of next-generation sequencing methods holds promise for yielding valuable insights into replication origin activation and their mapping in the mammalian genome, a fundamental yet largely enigmatic inquiry.

## SIGNIFICANCE

In previous research, the HaloTag-based sequential labeling technique employing one or two fluorescent ligands proved invaluable in uncovering distinct protein variants within histone, cohesin, and MCM complexes. These discoveries have advanced our understanding of the molecular pathways involved in maintaining genome stability, with direct implications for elucidating disease pathogenesis. Following the imaging studies, the next essential step requires the biochemical isolation of the distinct protein variants to facilitate a comprehensive examination of their biological properties. While the experimental HaloTag approach based on biotin-streptavidin interaction is well-suited for such studies, the practical implementation has proven to be challenging. In this work, we combined chemical synthesis, computer-assisted structural visualizations, and comprehensive cell biology-based characterization to examine several new Biotin-HaloTag ligands. Our results underscore the importance of the atom spacer (linker) length in effectively fine-tuning the HaloTag ligands and developing superior variants. The effort led to a new cell-permeable Biotin-[7]-HaloTag ligand 5 (Ligand 5), which exhibits efficient labeling of Halo-tagged proteins in the cellular environment of live cells and enables their pull-down through streptavidin beads without inducing cytotoxicity. Ligand 5 is demonstrated to effectively fill the gap in the spectrum of available HaloTag ligands for efficient cellular labeling and affinity pull-down of Halo-tagged proteins. Most notably, Ligand 5 will enable researchers to apply the HaloTag technology to studies of functional differences among protein variants using various biochemical approaches. Our work outlines the design of a sequential labeling scheme to effectively utilize the herein-developed Biotin-HaloTag ligand for the biochemical separation and detailed investigation of the biological properties of distinct protein variants. This encompasses the examination of interacting partners, post-translational modifications, and chromatin occupancy using advanced proteomic and next-generation sequencing techniques. Overall, we anticipate that this methodology will emerge as a robust approach for gaining mechanistic insights into fundamental cellular processes essential for organismal fitness and survival.

## ACKNOWLEDGMENTS

The research work in the Sedlackova laboratory was supported by the Czech Science Foundation Junior Star (grant no. 22-20303M), the European Union’s Horizon 2022 Widera Talent program (ERA grant agreement no. 101090292), EMBO Installation Grant (grant no. IG-5689-2024) and Jihomoravske centrum pro mezinarodni mobilitu (JCMM) project scholarship for foreign students. The research work in the Svenda laboratory was supported by the Bader Philanthropies and the National Infrastructure for Chemical Biology (CZ-OPENSCREEN, LM2023052). Vacha laboratory acknowledges funding from the project National Institute of Virology and Bacteriology (Programme EXCELES, ID Project No. LX22NPO5103) - Funded by the European Union - Next Generation EU.

We sincerely thank Jiri Polasek, Tomas Pop, and Tomas Jendrulek for the technical maintenance of high-content imaging microscopes. We are also deeply grateful to Maj-Britt Rask and Claudia Lukas for their support and for generously providing essential reagents. The HeLa Kyoto cell line was a gift from S. Narumiya. JFX554-HaloTag ligand was a kind gift from Luke Lavis. Special thanks to Claudia Lukas and Yimon Aye for their critical reading of the manuscript. We also express our gratitude to all members of the Sedlackova and Svenda laboratories for their stimulating discussions and insightful comments on the manuscript.

## AUTHOR CONTRIBUTION

H.P.-S. and J.S. conceived the project. A.K.Y. performed the majority of cell biology experiments, analyzed the data, and contributed to experimental design and concept development. A.J. and P.S. synthesized the Biotin-HaloTag ligands 3-6. P.F. performed some high-content imaging experiments, analyzed data, and contributed to concept development. I. K. and R. V. devised the structural visualization model and performed the calculations. H.P.-S. devised the original concept, generated cell lines, designed cell biology experiments, performed some labeling kinetics experiments, analyzed data, and prepared figures. H.P.-S. and J.S. wrote the manuscript. All authors read and commented on the manuscript.

## MATERIAL & CORRESPONDENCE

Should be addressed to J.S. or H.P.-S.

## METHODS

### Chemical synthesis of Biotin-HaloTag ligands

A detailed description of the experimental procedure for the synthesis of individual Biotin-HaloTag ligands and their structural analysis can be found in the Supplementary Information File 2.

### Computational model

The ternary complexes were assembled from the binary complexes of HaloTag-chloroalkane (PDB: 6U32, modified), streptavidin-biotin (PDB: 3RY2, one subunit of the tetramer), and a bifunctional ligand. A hydrocarbon spacer of varying lengths (0, 4, 7, 10, or 16 heavy atoms) was attached to the carboxyl group of biotin within the streptavidin-biotin complex. Exhaustive generation of linker conformers was performed using constrained embedding in RDKit (version 2023.03.3). Only the conformers with energy lower than 1 kcal/mol after 1000 steps of energy minimization, calculated using the MMFF94 force field, were retained. The RMSD threshold for pruning was set to 0.2 Å. The HaloTag-chloroalkane complex was then attached to the carboxyl group of each spacer conformation using the least squares fit of three atoms. The final ternary complexes were ranked by the number of overlapping heavy atoms, where two heavy atoms were considered overlapping if the distance between them was less than 3 Å.

### Cell culture

The human osteosarcoma cell line U2OS (ATCC HTB-96), the adenocarcinoma mammary epithelial cell line MCF7 (ATCC HTB-22), and the cervical HeLa Kyoto cell line (obtained from S. Narumiya) used in this study were cultured in Dulbecco’s modified Eagle’s medium (DMEM) containing high glucose and GlutaMAX (Thermo Fischer Scientific, 31966047) and supplemented with 10% Fetal Bovine Serum (FBS; Thermo Fischer Scientific, A5256801) along with 0.5% penicillin-streptomycin (Thermo Fischer Scientific, 15140122). The cells were grown under standard sterile conditions at 37°C and 5% CO_2_ level. CRISPR-Cas-9-mediated derivatives of U2OS cells expressing C-terminal endogenously tagged MCM2-Halo and MCM4-Halo were generated and validated in the previous study^23^. Supplementary Figure 1 and Supplementary Figure 5, respectively, provide additional validation of these cell lines.

To generate a cell line stably expressing H2B-Halo, the HeLa cells were transfected with a plasmid (pHCT-CMV-neo-H2B-Halo)^46^ using Lipofectamine LTX with Plus reagent (Thermo Fisher Scientific, 15338100). The transfected cells were selected with DMEM containing geneticin (G-418) (0.4 mg ml^-1^, Sigma-Aldrich, 4727878001) for 7 days, serially diluted and seeded onto 100 mm dishes, and grown under G-418 selection for an additional 12 days. The isolated colonies were expanded and tested for H2B-Halo expression by labeling with fluorescent JFX554 HaloTag ligand (labeling protocol is described in the section *HaloTag ligands and labeling protocol*). The generated H2B-Halo HeLa cell line was validated via SDS-PAGE and QIBC. All cell lines and their derivatives were regularly tested for mycoplasma using Mycoplasma Detection Kit (InvivoGen, rep-mys-50) and always found negative.

### Chemical reagents and antibodies

Verapamil (RayBiotech, 331-20649-1) was used as indicated in Figure panels. For the detection of BIOTIN-HaloTag ligand-tagged proteins, HRP-conjugated streptavidin (Thermo Fischer Scientific, 89880D, 1:1000) or Alexa Fluor 647-conjugated streptavidin (Thermo Fischer Scientific, S32357, 1:500) was used. Primary antibodies used for immunofluorescence (IF) were as follows: MCM2 (rabbit, Proteintech, 10513-1-AP, 1:1000), MCM4 (rabbit, Proteintech, 13043-1-AP, 1:1000) and PCNA (human, Immuno Concepts, 2037, 1:1000). Primary antibodies for western blotting (WB): ɑ-tubulin (rabbit, Proteintech, 11224-1-AP, 1:3000), Halo (mouse, Proteintech, 28A8, 1:1000), H3 (rabbit, Proteintech, 17168-1-AP, 1:5000), MCM2 (mouse, Novus Biologicals, H00004171-M01, 1:1000), MCM4 (rabbit, Proteintech, 13043-1-AP, 1:2000) and MCM7 (mouse, Santa Cruz, sc-9966, 1:1000). For IF, secondary antibody conjugates were goat anti-rabbit Alexa Fluor 647 (Thermo Fischer Scientific, A-21245, 1:2000) and donkey anti-human Alexa Fluor 647 (Jackson Immuno Research, 709-605-149, 1:2000). For WB, secondary antibody conjugates were HRP horse anti-mouse IgG antibody (Vector Laboratories, PI-2000, 1:10000) and horseradish peroxidase (HRP) goat anti-rabbit IgG antibody (Vector Laboratories, PI-1000-1, 1:10000).

### HaloTag ligands and labeling protocol

For single labeling by BIOTIN-HaloTag ligands, MCM4-Halo U2OS cells were incubated with indicated ligands at a final labeling concentration of 2.5 μM for 2 hours or increased concentrations as specified in figure panels for 2 hours. For concentration-dependent experiments, increasing ligand concentration was used as specified in figure panels for 2 hours. For all dual-HaloTag labeling, MCM4-Halo U2OS, MCM2-Halo U2OS, or H2B-Halo HeLa cells were pulsed with indicated BIOTIN-HaloTag ligands a final concentration of 2.5 μM for 2 hours or for the indicated time points as specified in figure panels. The BIOTIN-HaloTag ligands were then removed by washing cells three times with phosphate-buffered saline (PBS) buffer (Thermo Fisher Scientific, 14190250), and the cells were incubated with the JFX554-HaloTag ligand in a final concentration of 25 nM for 30 min based on kinetics experiments. For all single JFX554-HaloTag labeling, MCM4-Halo U2OS, MCM2-Halo U2OS, or H2B-Halo HeLa cells were incubated with the JFX554-HaloTag ligand in a final concentration of 25 nM for 30 min. After HaloTag labeling, cell samples were further processed for western blotting or IF staining.

### SDS-PAGE and western blotting

Whole-cell lysates were obtained by incubating cells in lysis buffer (10 mM Tris-Cl pH 7.5, 150 mM NaCl, 0.5 mM EDTA, 0.5% NP-40) supplemented with protease and phosphatase inhibitors (ROCHE, 4693116001 and 4906845001) and 250 U per ml of benzonase (Merck, E1014-25KU) on ice for 45 min. For the preparation of whole cell lysates from H2B-Halo HeLa cells, cell lysis on ice was followed by sonication. The cell lysates were denatured at 95°C for 5 min in Laemmli loading buffer containing NuPAGE LDS Sample Buffer (Thermo Fisher Scientific, NP0007) and NuPAGE Sample Reducing Agent (Thermo Fisher Scientific, NP0009) and separated on mPAGE™ 4-12% Bis-Tris Precast Gels (Merck, MP41G10 or MP41G12) through standard SDS-PAGE protocol. After SDS-PAGE, gels were scanned with Cy3 wavelength to detect MCM4-Halo or H2B-Halo labeled by fluorescent JFX554 HaloTag ligand. Afterward, the separated proteins were transferred from gel to nitrocellulose membrane (Thermo Fisher Scientific, IB23002X3) using iBlot (Thermo Fisher Scientific, IB21001). Ponceau S solution (Merck, P7170-1L) was used for total protein staining. To detect MCM4-Halo, MCM2-Halo, or H2B-Halo labeled by non-fluorescent Biotin-HaloTag ligands, the membrane was blocked with Nucleic Acid Detection Blocking Buffer (NADB) buffer (Thermo Fischer Scientific, 89880A) for 30 min followed by incubation with HRP-coupled streptavidin (Thermo Fischer Scientific, 89880D) in NADB buffer for 1 hour at room temperature. To detect the target proteins, the membrane was blocked with PBS buffer containing 5% milk (Merck, 70166-500G) and 0.1% Tween-20 (Merck, P1379-500ML) for 1 hour at room temperature, followed by primary and secondary antibody staining. Both primary and secondary antibodies were diluted in PBS buffer containing 5% milk and 0.1% Tween-20 and then incubated overnight at 4°C or 2 hours at room temperature, respectively. After incubation, HRP-signal was detected by ECL Select Western Blotting Detection Reagent (Merck, GERPN2235). After the detection of target proteins, the membrane was stripped using Restore PLUS Western Blot Stripping Buffer (Thermo Fischer Scientific, 46430) and stained with MCM7 (Supplementary Figure 1E), ɑ-tubulin (Supplementary Figures 1A, 5E) or H3 (Figure 4E) antibodies as loading controls. For processing controls (Figures 1C, 2F, 3C; Supplementary Figure 3A), the same lysates and loading concentrations were used, and the samples were run parallelly to detect target proteins.

### Streptavidin-based pull-down

Whole-cell lysates obtained from indicated cells were incubated with either 40 µL of anti-streptavidin agarose beads (Thermo Fisher Scientific, 20347) for 2 hours at 4°C (Figures 1F, 2H, 3G, 4D, 4F, Supplementary Figure 4G) or 100 µL of anti-streptavidin agarose beads overnight at 4°C (Figure 3H). Beads were then washed twice with low salt wash buffer (10 mM Tris-Cl pH 7.5, 150 mM NaCl, 0.5 mM EDTA, 0.5% NP-40) supplemented with protease and phosphatase inhibitors and twice with high salt wash buffer (10 mM Tris-Cl pH 7.5, 500 mM NaCl, 0.5 mM EDTA, 0.5% NP-40) supplemented with protease and phosphatase inhibitors. To elute bound proteins, beads were incubated with 60 µL of Laemmli loading buffer for 20 min at 95°C. The elutes were analyzed by western blotting using either streptavidin-HRP to detect BIOTIN-HaloTag ligand-labeled proteins or specific antibodies, as indicated in the figures.

### IF staining

For IF staining, cells were grown on round 12-mm diameter, 1.5-mm-thick glass coverslips (VWR, MENZCB00120RAC20). For staining of chromatin-bound proteins, cells were pre-extracted with ice-cold cytoskeleton buffer (10 mM Hepes pH 7.5; 300 mM sucrose; 100 mM NaCl; 3 mM MgCl_2_ and 0.5% Triton X-100) for 10 min at room temperature, washed three times with PBS and fixed using 4% buffered formaldehyde (VWR, 9713.1000) for 20 min at room temperature. To stain total nuclear proteins, cells were fixed directly by 4% buffered formaldehyde for 20 min at room temperature, washed three times with PBS, and incubated with ice-cold PBS containing 0.2% TritonX-100 for 5 min at room temperature. Both primary and secondary antibodies were diluted in fresh DMEM supplemented with 10% FBS and incubated with coverslips at room temperature for 90 min and 45 min, respectively. To counterstain nuclear DNA, the secondary antibody cocktail was supplemented with 0.5 µg per ml 4’,6-Diamidino-2-Phenylindole, Dihydrochloride (DAPI; Thermo Fisher Scientific, D1306). After secondary antibody incubation, the coverslips were washed three times with PBS and twice with distilled water, air dried, and mounted on slides using Mowiol-based mounting medium (12% Mowiol 4-88 (Merck, 81381-250G), 30% glycerol, 0.12 M Tris-HCl pH 8.5). To detect Halo-tagged proteins labeled by Biotin-HaloTag ligands, cells were fixed or pre-extracted as described above. Coverslips were blocked with Nucleic Acid Detection Blocking Buffer buffer (Thermo Fischer Scientific, 89880A) for 30 min at room temperature, washed three times with PBS, and incubated with Alexa Fluor 647-conjugated streptavidin (Thermo Fischer Scientific, S32357) and DAPI in PBS buffer for 1 hour at room temperature. After incubation, the coverslips were processed as described above.

### Quantitative Image-Based cytometry

Images were acquired using an inverted screening microscope ScanR (IX83, Evident), equipped with a UPLXAPO dry objective (20X, 0.8 NA); fast excitation and emission filter wheel for DAPI, FITC, Cy3, and Cy5 wavelengths, Lumencor Spectra X led fluorescence light source and digital monochromatic sCMOS ORCA-Flash 4.0 LT Plus camera. The images were acquired with the ScanR acquisition software (Evident, v.3.4.1) in an automated fashion with constant laser intensity at 100% and exposure times adjusted individually for each fluorophore. The automated image analysis was subsequently performed in ScanR analysis software (Evident, v.3.4.1). Automated dynamic background correction thresholding at least fivefold pixel intensity above background levels was applied for each fluorescent channel separately to maintain the same conditions within a single experiment. An intensity-threshold-based mask was generated based on the DAPI signal to identify individual nuclei as main objects. This mask was then applied to analyze pixel intensities in different channels for each nucleus. The multiparameter analysis was exported as a table and further analyzed in Spotfire software (TIBCO, v.11.8.0). A similar number of cells was analyzed and compared within each experiment.

### Colony formation assay

U2OS or MCF7 cells were seeded onto 6-well plates (400 cells per well) containing complete DMEM supplemented with increasing concentration of verapamil or Ligand 5. After ten days, cells were fixed using 4% buffered formaldehyde for 20 min at room temperature and stained with 0.1% crystal violet diluted in 20% ethanol for 30 min at room temperature. The plates were washed and air-dried overnight before counting the colonies.

## REFERENCES

1 Terpe, K. Overview of tag protein fusions: from molecular and biochemical fundamentals to commercial systems. Appl Microbiol Biotechnol 60, 523–533, doi:10.1007/s00253-002-1158-6 (2003).

2 Giepmans, B. N., Adams, S. R., Ellisman, M. H. & Tsien, R. Y. The fluorescent toolbox for assessing protein location and function. Science 312, 217–224, doi:10.1126/science.1124618 (2006).

3 Yesbolatova, A. et al. The auxin-inducible degron 2 technology provides sharp degradation control in yeast, mammalian cells, and mice. Nat Commun 11, 5701, doi:10.1038/s41467-020-19532-z (2020).

4 Bond, A. G. et al. Development of BromoTag: A “Bump-and-Hole”-PROTAC System to Induce Potent, Rapid, and Selective Degradation of Tagged Target Proteins. J Med Chem 64, 15477–15502, doi:10.1021/acs.jmedchem.1c01532 (2021).

5 Nabet, B. et al. The dTAG system for immediate and target-specific protein degradation. Nat Chem Biol 14, 431–441, doi:10.1038/s41589-018-0021-8 (2018).

6 Keppler, A. et al. A general method for the covalent labeling of fusion proteins with small molecules in vivo. Nat Biotechnol 21, 86–89, doi:10.1038/nbt765 (2003).

7 Los, G. V. et al. HaloTag: a novel protein labeling technology for cell imaging and protein analysis. ACS Chem Biol 3, 373–382, doi:10.1021/cb800025k (2008).

8 Grimm, J. B. et al. A general method to optimize and functionalize red-shifted rhodamine dyes. Nat Methods 17, 815–821, doi:10.1038/s41592-020-0909-6 (2020).

9 Deo, C. et al. The HaloTag as a general scaffold for far-red tunable chemigenetic indicators. Nat Chem Biol 17, 718–723, doi:10.1038/s41589-021-00775-w (2021).

10 Buckley, D. L. et al. HaloPROTACS: Use of Small Molecule PROTACs to Induce Degradation of HaloTag Fusion Proteins. ACS Chem Biol 10, 1831–1837, doi:10.1021/acschembio.5b00442 (2015).

11 Wang, L. et al. A general strategy to develop cell permeable and fluorogenic probes for multicolour nanoscopy. Nat Chem 12, 165–172, doi:10.1038/s41557-019-0371-1 (2020).

12 Frei, M. S. et al. Engineered HaloTag variants for fluorescence lifetime multiplexing. Nat Methods 19, 65–70, doi:10.1038/s41592-021-01341-x (2022).

13 Murrey, H. E. et al. Systematic Evaluation of Bioorthogonal Reactions in Live Cells with Clickable HaloTag Ligands: Implications for Intracellular Imaging. J Am Chem Soc 137, 11461–11475, doi:10.1021/jacs.5b06847 (2015).

14 Torne, J., Orsi, G. A., Ray-Gallet, D. & Almouzni, G. Imaging Newly Synthesized and Old Histone Variant Dynamics Dependent on Chaperones Using the SNAP-Tag System. Methods Mol Biol 1832, 207–221, doi:10.1007/978-1-4939-8663-7_11 (2018).

15 Adam, S. et al. Real-Time Tracking of Parental Histones Reveals Their Contribution to Chromatin Integrity Following DNA Damage. Mol Cell 64, 65–78, doi:10.1016/j.molcel.2016.08.019 (2016).

16 Stewart-Morgan, K. R., Petryk, N. & Groth, A. Chromatin replication and epigenetic cell memory. Nat Cell Biol 22, 361–371, doi:10.1038/s41556-020-0487-y (2020).

17 Torne, J. et al. Two HIRA-dependent pathways mediate H3.3 de novo deposition and recycling during transcription. Nat Struct Mol Biol 27, 1057–1068, doi:10.1038/s41594-020-0492-7 (2020).

18 Adam, S., Polo, S. E. & Almouzni, G. Transcription recovery after DNA damage requires chromatin priming by the H3.3 histone chaperone HIRA. Cell 155, 94–106, doi:10.1016/j.cell.2013.08.029 (2013).

19 Jansen, L. E., Black, B. E., Foltz, D. R. & Cleveland, D. W. Propagation of centromeric chromatin requires exit from mitosis. J Cell Biol 176, 795–805, doi:10.1083/jcb.200701066 (2007).

20 Saredi, G. et al. The histone chaperone SPT2 regulates chromatin structure and function in Metazoa. Nat Struct Mol Biol 31, 523–535, doi:10.1038/s41594-023-01204-3 (2024).

21 Srinivasan, M., Fumasoni, M., Petela, N. J., Murray, A. & Nasmyth, K. A. Cohesion is established during DNA replication utilising chromosome associated cohesin rings as well as those loaded de novo onto nascent DNAs. Elife 9, doi:10.7554/eLife.56611 (2020).

22 Rhodes, J. D. P. et al. Cohesin Can Remain Associated with Chromosomes during DNA Replication. Cell Rep 20, 2749–2755, doi:10.1016/j.celrep.2017.08.092 (2017).

23 Sedlackova, H. et al. Equilibrium between nascent and parental MCM proteins protects replicating genomes. Nature 587, 297–302, doi:10.1038/s41586-020-2842-3 (2020).

24 Yatskevich, S., Rhodes, J. & Nasmyth, K. Organization of Chromosomal DNA by SMC Complexes. Annu Rev Genet 53, 445–482, doi:10.1146/annurev-genet-112618-043633 (2019).

25 Alonso-Gil, D. & Losada, A. NIPBL and cohesin: new take on a classic tale. Trends Cell Biol 33, 860–871, doi:10.1016/j.tcb.2023.03.006 (2023).

26 Yadav, A. K. & Polasek-Sedlackova, H. Quantity and quality of minichromosome maintenance protein complexes couple replication licensing to genome integrity. Commun Biol 7, 167, doi:10.1038/s42003-024-05855-w (2024).

27 Pratik Kumar, J. D. V., Edwin R. Chapman, Luke D. Lavis. Multifunctional fluorophores for live-cell imaging and affinity capture of proteins. bioRxiv, 10.1101/2022.07.02.498544 (2022).

28 Promega. HaloTag® PEG-Biotin Ligand Protocol (technical note # 9PIG859); https://worldwide.promega.com/resources/protocols/product-information-sheets/g/halotag-pegbiotin-ligand-protocol/. (2016).

29 Payne, N. C., Kalyakina, A. S., Singh, K., Tye, M. A. & Mazitschek, R. Bright and stable luminescent probes for target engagement profiling in live cells. Nat Chem Biol 17, 1168–1177, doi:10.1038/s41589-021-00877-5 (2021).

30 Sridharan, S. et al. Systematic discovery of biomolecular condensate-specific protein phosphorylation. Nat Chem Biol 18, 1104–1114, doi:10.1038/s41589-022-01062-y (2022).

31 Lukinavicius, G. et al. Fluorogenic probes for live-cell imaging of the cytoskeleton. Nat Methods 11, 731–733, doi:10.1038/nmeth.2972 (2014).

32 Kania, E. et al. Verapamil treatment induces cytoprotective autophagy by modulating cellular metabolism. FEBS J 284, 1370–1387, doi:10.1111/febs.14064 (2017).

33 So, M. K., Yao, H. & Rao, J. HaloTag protein-mediated specific labeling of living cells with quantum dots. Biochem Biophys Res Commun 374, 419–423, doi:10.1016/j.bbrc.2008.07.004 (2008).

34 Klein, V. G., Bond, A. G., Craigon, C., Lokey, R. S. & Ciulli, A. Amide-to-Ester Substitution as a Strategy for Optimizing PROTAC Permeability and Cellular Activity. J Med Chem 64, 18082–18101, doi:10.1021/acs.jmedchem.1c01496 (2021).

35 Le Trong, I. et al. Streptavidin and its biotin complex at atomic resolution. Acta Crystallogr D Biol Crystallogr 67, 813–821, doi:10.1107/S0907444911027806 (2011).

36 Polasek-Sedlackova, H., Miller, T. C. R., Krejci, J., Rask, M. B. & Lukas, J. Solving the MCM paradox by visualizing the scaffold of CMG helicase at active replisomes. Nat Commun 13, 6090, doi:10.1038/s41467-022-33887-5 (2022).

37 Saxton, M. N., Morisaki, T., Krapf, D., Kimura, H. & Stasevich, T. J. Live-cell imaging uncovers the relationship between histone acetylation, transcription initiation, and nucleosome mobility. Sci Adv 9, eadh4819, doi:10.1126/sciadv.adh4819 (2023).

38 England, C. G., Luo, H. & Cai, W. HaloTag technology: a versatile platform for biomedical applications. Bioconjug Chem 26, 975–986, doi:10.1021/acs.bioconjchem.5b00191 (2015).

39 Grimm, J. B. et al. A General Method to Improve Fluorophores Using Deuterated Auxochromes. JACS Au 1, 690–696, doi:10.1021/jacsau.1c00006 (2021).

40 Kompa, J. et al. Exchangeable HaloTag Ligands for Super-Resolution Fluorescence Microscopy. J Am Chem Soc 145, 3075–3083, doi:10.1021/jacs.2c11969 (2023).

41 Petryk, N. et al. MCM2 promotes symmetric inheritance of modified histones during DNA replication. Science 361, 1389–1392, doi:10.1126/science.aau0294 (2018).

42 Yu, C. et al. A mechanism for preventing asymmetric histone segregation onto replicating DNA strands. Science 361, 1386–1389, doi:10.1126/science.aat8849 (2018).

43 Li, Z. et al. DNA polymerase alpha interacts with H3-H4 and facilitates the transfer of parental histones to lagging strands. Sci Adv 6, eabb5820, doi:10.1126/sciadv.abb5820 (2020).

44 Flury, V. et al. Recycling of modified H2A-H2B provides short-term memory of chromatin states. Cell 186, 1050–1065 e1019, doi:10.1016/j.cell.2023.01.007 (2023).

45 Charlton, S. J. et al. The fork protection complex promotes parental histone recycling and epigenetic memory. Cell, doi:10.1016/j.cell.2024.07.017 (2024).

46 Ochs, F. et al. Stabilization of chromatin topology safeguards genome integrity. Nature 574, 571–574, doi:10.1038/s41586-019-1659-4 (2019).

