## Supplementary Figures 1-6 for "Development of a cell-permeable Biotin-HaloTag ligand to explore functional differences between protein variants across cellular generations"

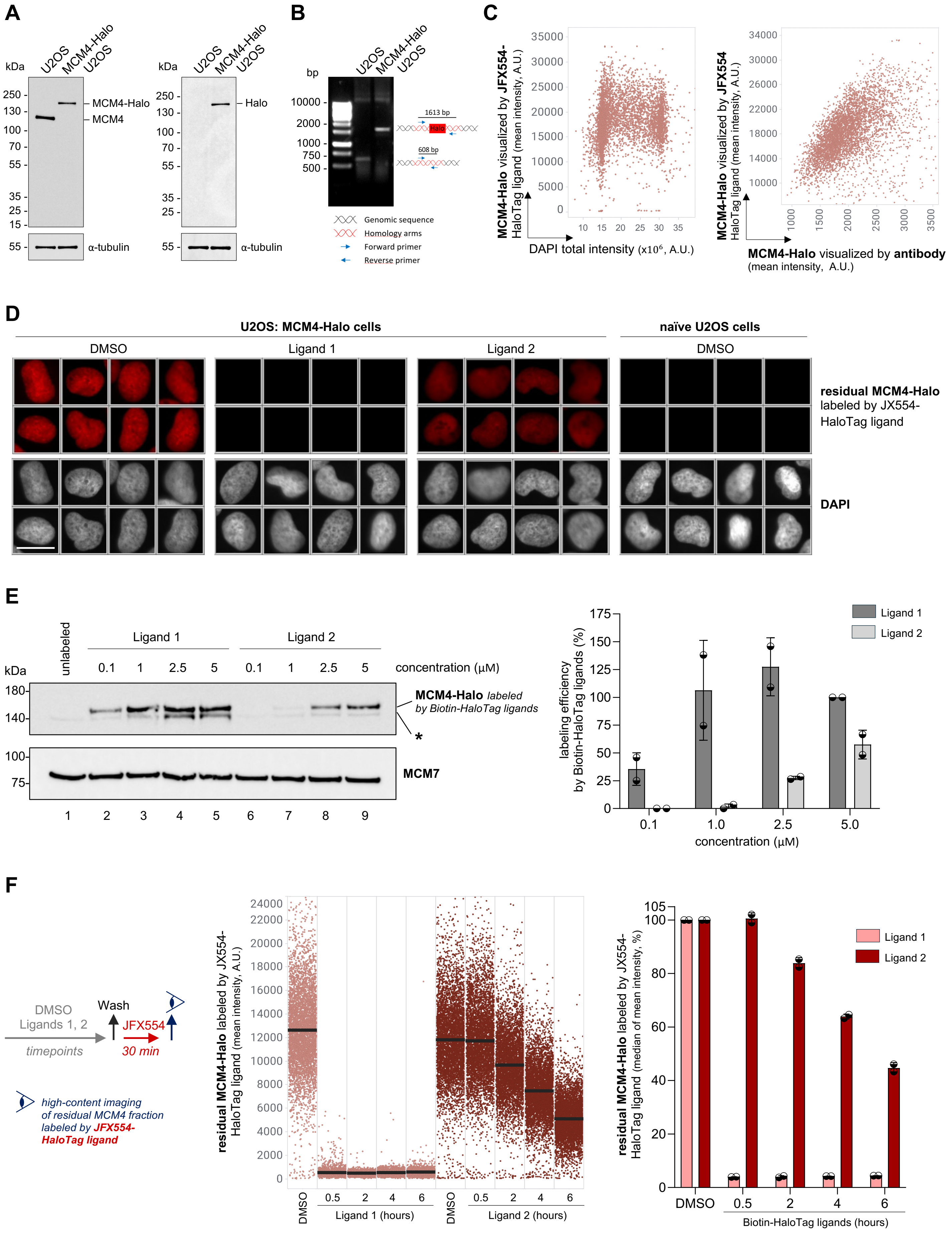


**Supplementary Figure 1**

**Supplementary Figure 1: Validation of MCM4-Halo cell line and assessment of labeling kinetics of commercial Biotin-HaloTag ligands. (A)** Western blots of U2OS and MCM4-Halo U2OS cells stained for MCM4 (left) or Halo (right). α-tubulin was used as a loading control. **(B)** Junction PCR showing homozygous MCM4-Halo tagging. **(C)** QIBC of MCM4-Halo U2OS cells pulsed with JFX554-HaloTag ligand for 30 min and immunostained for MCM4. Nuclear DNA was counterstained with DAPI (*n* ≈ 5000 cells per condition). **(D)** Unbiased QIBC galleries of residual MCM4-Halo labeled by JFX554-HaloTag ligand following labeling by indicated Biotin-HaloTag ligands. Nuclear DNA was counterstained with DAPI. See the pulse-chase protocol and QIBC analysis in Figure 1E. Scale bar, 20 µm. **(E)** Left, western blotting of whole cell lysates of MCM4-Halo U2OS cells labeled with Biotin-HaloTag ligands with increasing concentration as indicated for 2 hours. MCM7 was stained as a loading control. Right, quantification of labeling efficiency for indicated Biotin-HaloTag ligands based on western blot on left. **(F)** Left, the pulse-chase protocol of MCM4-Halo U2OS cells labeled with indicated HaloTag ligands. Middle, QIBC of the residual fraction of MCM4-Halo labeled by JFX554-HaloTag ligand. Nuclear DNA was counterstained with DAPI. Lines denote medians; *n* ≈ 5000 cells per condition. Right, the quantification of the QIBC plot in the middle. Each bar indicates the median of mean intensity normalized with respect to DMSO as 100 percent; *n* = 2 technical replicates.

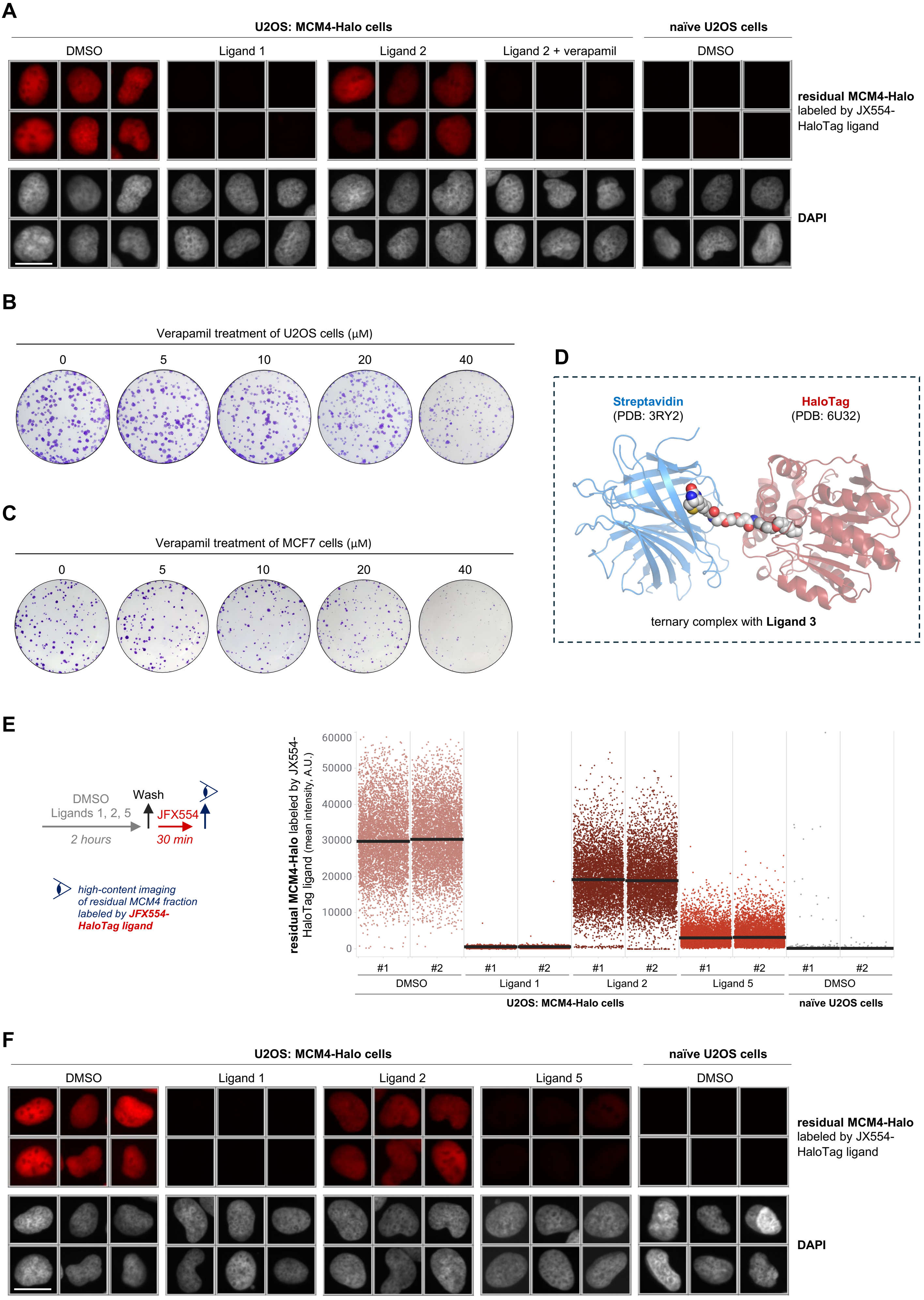


### Supplementary Figure 2

**Supplementary Figure 2: Ligand 5 effectively labels the Halo-tagged MCM4 protein subunit in the cellular environment. (A)** Unbiased QIBC galleries of residual MCM4-Halo labeled by JFX554-HaloTag ligand following labeling by indicated Biotin-HaloTag ligands with or without verapamil. Nuclear DNA was counterstained with DAPI. See the pulse-chase protocol and QIBC analysis in Figure 2B. Scale bar, 20 µm. **(B)** Representative images of colonies formed after verapamil treatment of U2OS cells at indicated concentrations. See quantification in Figure 2C. **(C)** Representative images of colonies formed after verapamil treatment of MCF7 cells at indicated concentrations. See quantification in Figure 2C. **(D)** Representative structural visualization of the HaloTag-ligand-streptavidin ternary complex for Ligand 3. The complexes were calculated from the binary complexes of streptavidin-biotin (PDB: 3RY2, one subunit of the tetramer) and HaloTag-chloroalkane (PDB: 6U32, modified), with the exhaustive generation of spacer conformations by constrained embedding in RDKit. **(E)** Left, the pulse-chase protocol of MCM4-Halo U2OS cells labeled with indicated HaloTag ligands. Right, QIBC of the residual fraction of MCM4-Halo labeled by JFX554-HaloTag ligand. Nuclear DNA was counterstained with DAPI. Lines denote medians; $n \approx$ 6000 cells per condition. See quantification in Figure 3F. **(F)** Unbiased QIBC galleries of residual MCM4-Halo labeled by JFX554-HaloTag ligand following labeling by indicated Biotin-HaloTag ligands. Nuclear DNA was counterstained with DAPI. See the pulse-chase protocol and QIBC analysis in (D) and Figure 3F. Scale bar, 20 µm.

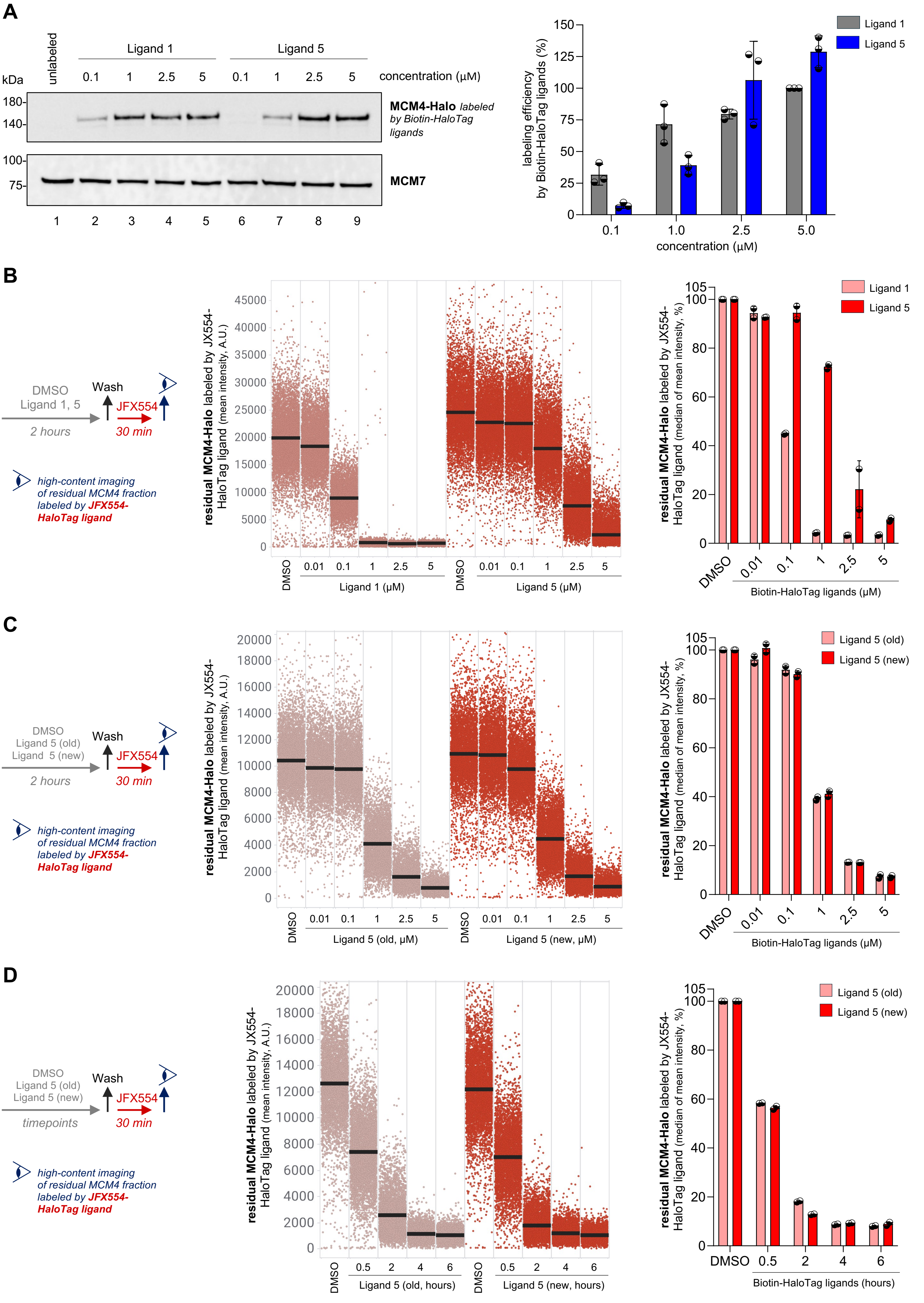


**Supplementary Figure 3**

**Supplementary Figure 3: Assessment of labeling kinetics of commercial Ligand 1 and two different batches of Ligand 5 in a concentration- and time-dependent manner. (A)** Left, western blotting of whole cell lysates of MCM4-Halo U2OS cells labeled with Biotin-HaloTag ligands with increasing concentration as indicated. MCM7 was used as a processing control. Right, quantification of labeling efficiency for indicated Biotin-HaloTag ligands based on western blot on left. **(B)** Left, the pulse-chase protocol of MCM4-Halo U2OS cells labeled with indicated HaloTag ligands. Middle, QIBC of the residual fraction of MCM4-Halo labeled by JFX554-HaloTag ligand. Nuclear DNA was counterstained with DAPI. Lines denote medians; *n* ≈ 7000 cells per condition. Right, the quantification of QIBC plots in the middle. Each bar indicates the median of mean intensity normalized with respect to DMSO as 100 percent; *n* = 2 technical replicates. **(C)** Left, the pulse-chase protocol of MCM4-Halo U2OS cells labeled with indicated HaloTag ligands. Middle, QIBC of the residual fraction of MCM4-Halo labeled by JFX554-HaloTag ligand. Nuclear DNA was counterstained with DAPI. Lines denote medians; *n* ≈ 5000 cells per condition. Right, the quantification of QIBC plots in the middle. Each bar indicates the median of mean intensity normalized with respect to DMSO as 100 percent; *n* = 2 technical replicates. **(D)** Left, the pulse-chase protocol of MCM4-Halo U2OS cells labeled with indicated HaloTag ligands. Middle, QIBC of the residual fraction of MCM4-Halo labeled by JFX554-HaloTag ligand. Nuclear DNA was counterstained with DAPI. Lines denote medians; *n* ≈ 5000 cells per condition. Right, the quantification of QIBC plots in the middle. Each bar indicates the median of mean intensity normalized with respect to DMSO as 100 percent; *n* = 2 technical replicates.

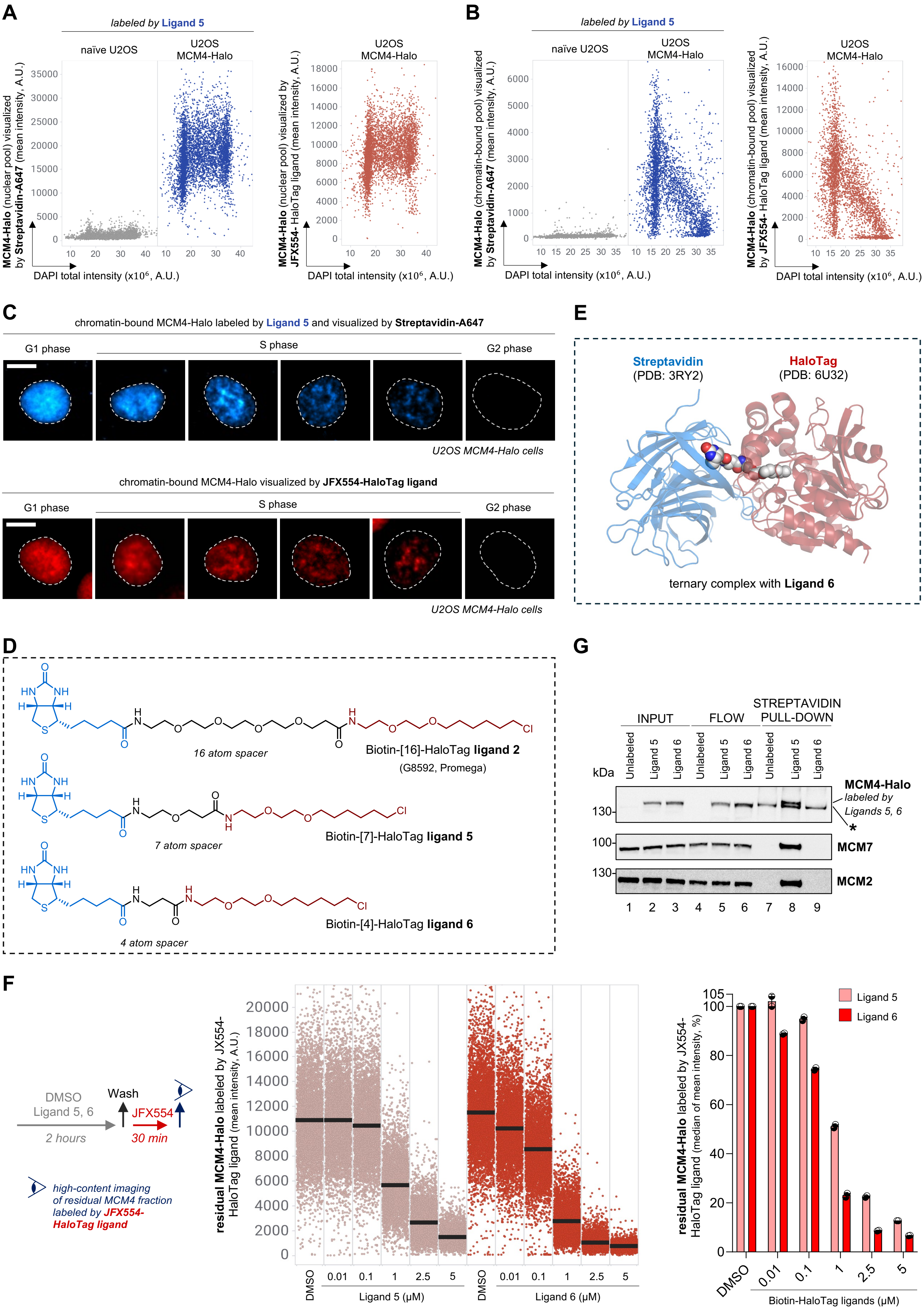


**Supplementary Figure 4**

**Supplementary Figure 4: Single-cell-based visualization of MCM complexes dynamics during cell cycle using Ligand 5 and fluorescent streptavidin. (A)** QIBC of a nuclear fraction of MCM4-Halo complexes in naïve and MCM4-Halo U2OS cells pulsed with Ligand 5 (left) or JFX554-HaloTag ligand (right) for 2 hours. Nuclear DNA was counterstained with DAPI (*n* ≈ 5000 cells per condition). **(B)** QIBC of chromatin-bound MCM4-Halo complexes in naïve and MCM4-Halo U2OS cells pulsed with Ligand 5 (left) or JFX554-HaloTag ligand (right) for 2 hours. Nuclear DNA was counterstained with DAPI (*n* ≈ 3500 cells per condition). **(C)** Representative widefield microscopy images capturing chromatin dynamics of MCM complexes visualized by Streptavidin-A647 (top) or JFX554-HaloTag ligand (bottom). Scale bar, 10 µm. **(D)** Chemical structure of commercially available Biotin-HaloTag ligand (top) with 16-atom spacer compared to newly designed Biotin-HaloTag ligands with 7-atom spacer (middle) and 4-atom spacer (bottom). **(E)** Representative structural visualization of the HaloTag-ligand-streptavidin ternary complex for Ligand 6. The complexes were calculated from the binary complexes of streptavidin-biotin (PDB: 3RY2, one subunit of the tetramer) and HaloTag-chloroalkane (PDB: 6U32, modified), with the exhaustive generation of spacer conformations by constrained embedding in RDKit. **(F)** Left, the pulse-chase protocol of MCM4-Halo U2OS cells labeled with indicated HaloTag ligands. Middle, QIBC of the residual fraction of MCM4-Halo labeled by JFX554-HaloTag ligand. Nuclear DNA was counterstained with DAPI. Lines denote medians; *n* ≈ 5000 cells per condition. Right, the quantification of QIBC plots in the middle. Each bar indicates the median of mean intensity normalized with respect to DMSO as 100 percent; *n* = 2 technical replicates. **(G)** Streptavidin pull-down of whole cell lysates of MCM4-Halo U2OS cells labeled with indicated Biotin-HaloTag ligands at a final concentration of 2.5 µM for 2 hours.

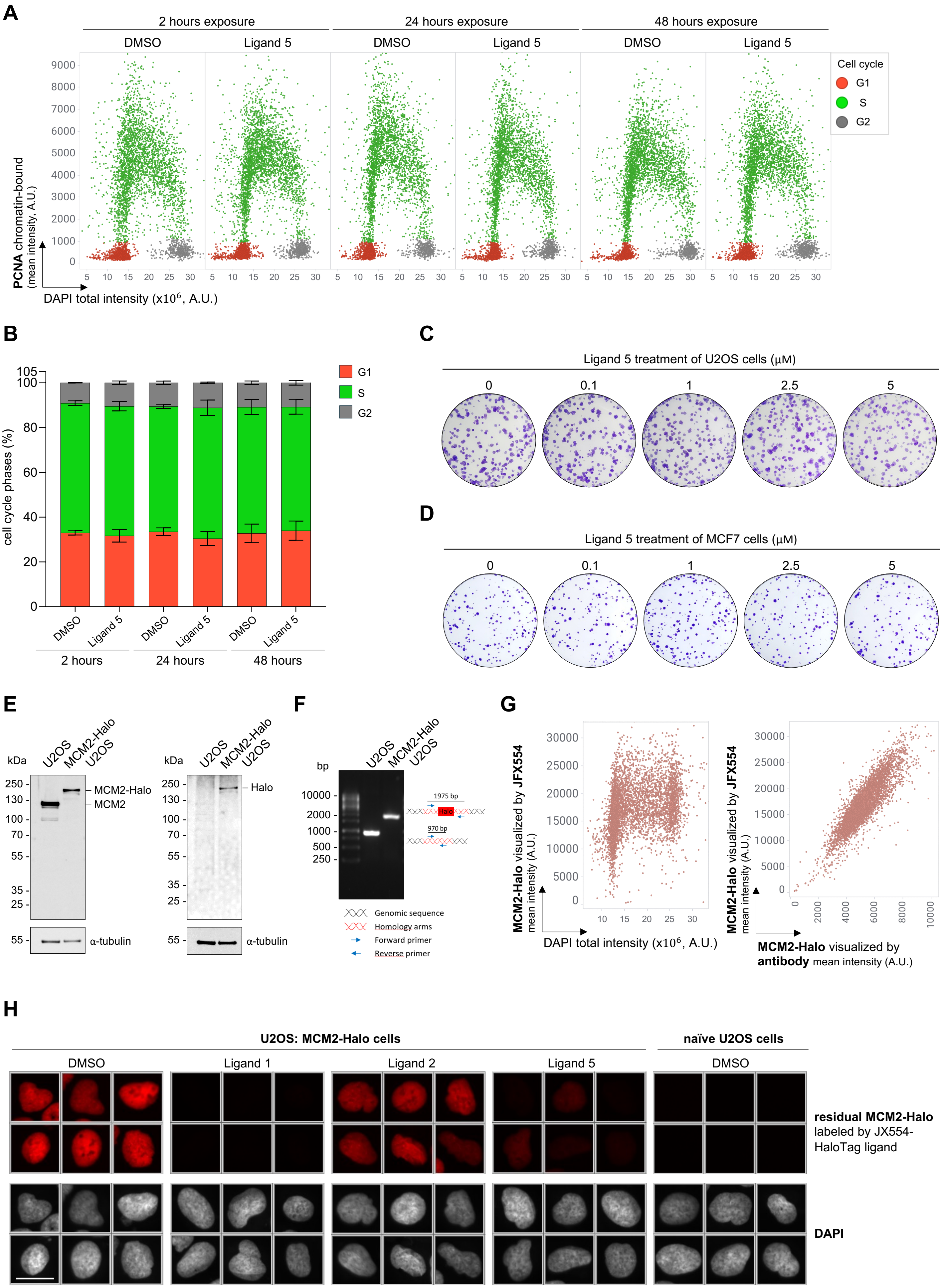


Supplementary Figure 5

**Supplementary Figure 5: Ligand 5 does not affect cell cycle or cell growth during long-term experiments. (A)** QIBC of chromatin-bound PCNA in MCM4-Halo U2OS cells upon treatment with Ligand 5 at a final concentration of 2.5 µM for indicated timepoints. Nuclear DNA was counterstained with DAPI; *n* ≈ 5000 cells per condition. **(B)** Quantification of individual cell cycle phases based on QIBC in (D); *n* = 2 technical replicates. **(C)** Representative images of colonies formed after Ligand 5 treatment of U2OS cells at indicated concentrations. See quantification in Figure 4A. **(D)** Representative images of colonies formed after Ligand 5 treatment of MCF7 cells at indicated concentrations. See quantification in Figure 4A. **(E)** Western blots of U2OS and MCM4-Halo U2OS cells stained for MCM4 (left) or Halo (right). α-tubulin was used as a loading control. **(F)** Junction PCR showing homozygous MCM2-Halo tagging. **(G)** QIBC of MCM2-Halo U2OS cells pulsed with JFX554 HaloTag ligand for 30 min and immunostained for MCM2. Nuclear DNA was counterstained with DAPI (*n* ≈ 5000 cells per condition). **(H)** Unbiased QIBC galleries of residual MCM2-Halo labeled by JFX554-HaloTag ligand following labeling by indicated Biotin-HaloTag ligands. Nuclear DNA was counterstained with DAPI. See the pulse-chase protocol and QIBC analysis in Figure 4C. Scale bar, 20 µm.

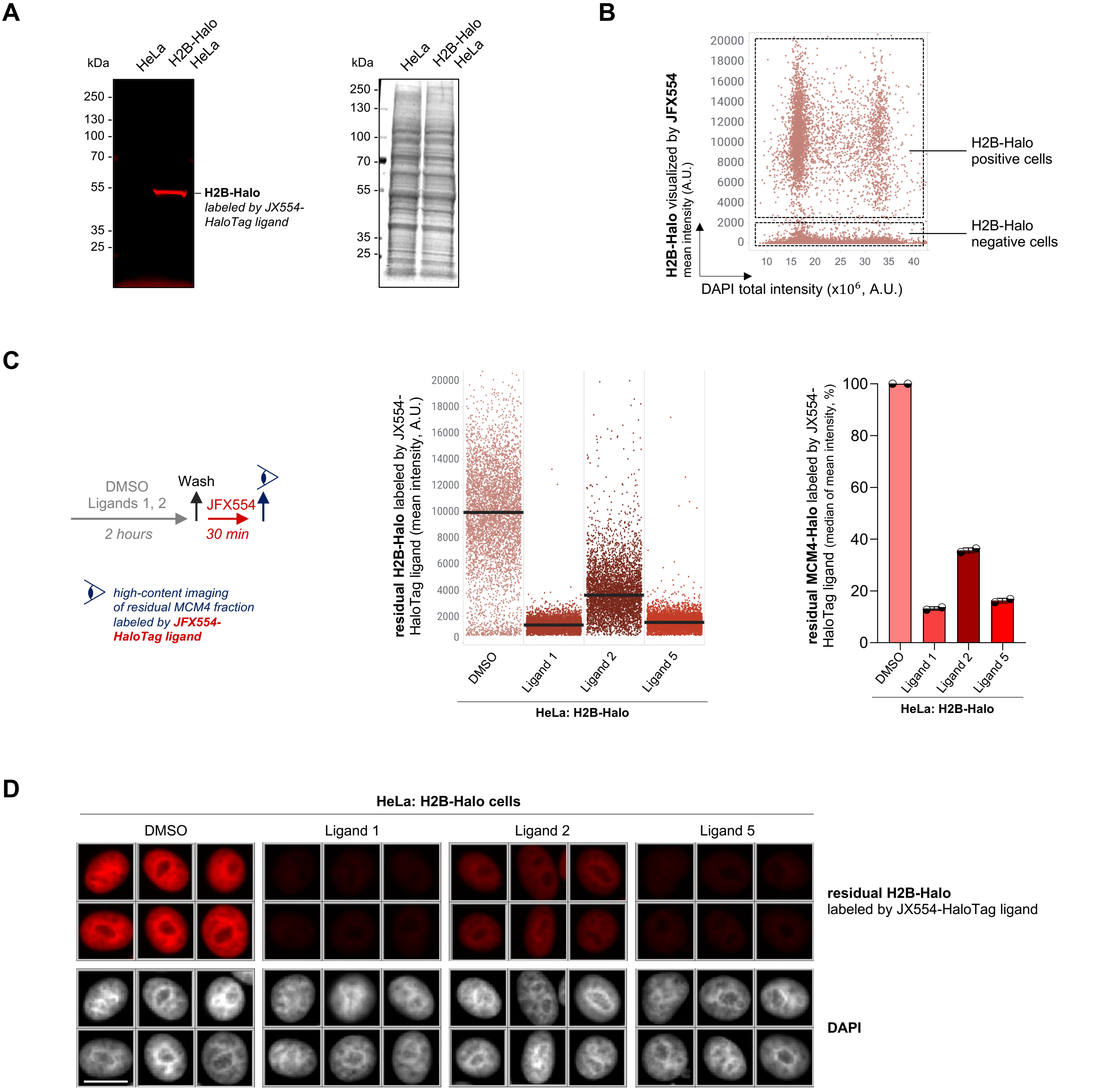


**Supplementary Figure 6**

**Supplementary Figure 6: Ligand 5 shows the high labeling efficiency of H2B-Halo in HeLa cells. (A)** Left, SDS-PAGE of whole cell lysates of HeLa and H2B-Halo HeLa cells labeled with JFX554-HaloTag ligand. Right, total protein staining. **(B)** QIBC of H2B-Halo HeLa cells pulsed with JFX554-HaloTag ligand for 30 min. Nuclear DNA was counterstained with DAPI (*n* ≈ 5000 cells). **(C)** Left, the pulse-chase protocol of H2B-Halo HeLa cells labeled with indicated HaloTag ligands. Middle, QIBC of the residual fraction of H2B-Halo labeled by JFX554-HaloTag ligand. Nuclear DNA was counterstained with DAPI. Lines denote medians; *n* ≈ 5000 cells per condition. Right, the quantification of QIBC plots in the middle. Each bar indicates the median of mean intensity normalized with respect to DMSO as 100 percent; *n* = 2 technical replicates. **(D)** Unbiased QIBC galleries of residual H2B-Halo labeled by JFX554-HaloTag ligand following labeling by indicated Biotin-HaloTag ligands. Nuclear DNA was counterstained with DAPI. See the pulse-chase protocol and QIBC analysis in (C). Scale bar, 20 µm.
